# Spatial metabolomics for symbiotic marine invertebrates

**DOI:** 10.1101/2022.10.05.511040

**Authors:** Wing Yan Chan, David Rudd, Madeleine J. H. van Oppen

## Abstract

Microbial symbionts frequently localize within specific body structures or cell types of their multicellular hosts. This spatiotemporal niche is critical to host health, nutrient exchange and fitness. Measuring host-microbe metabolite exchange has conventionally relied on tissue homogenates, eliminating dimensionality and dampening analytical sensitivity. We have developed a mass spectrometry imaging (MSI) workflow for a soft- and hard-bodied cnidarian animal capable of revealing the host and symbiont metabolome *in situ*, without the need for *a priori* isotopic labelling or skeleton decalcification. The MSI method provides critical functional insights that cannot be gleaned from bulk tissue analyses or other presently available spatial methods. We show that cnidarian hosts may regulate microalgal symbionts acquisition and rejection through specific ceramides distributed throughout the tissue lining the gastrovascular cavity; once resident, symbionts reside in light-exposed tentacles to generate photosynthate. These spatial patterns reveal how symbiont identity can drive host metabolism.

## INTRODUCTION

Microorganisms such as eukaryotic microalgae, bacteria, viruses, fungi and archaea are the most abundant life forms on Earth. Their symbioses with multicellular hosts can improve host uptake of nutrients (McFall-Ngai *et al,* 2013), vitamins and minerals; mitigate toxic compounds and control pathogens; play a critical role in host health (e.g., human gut microbiota); and form the basis of the ecological success of some of the most productive ecosystems on earth (e.g., coral reefs, tropical rainforests, deep sea hydrothermal vents) (Peixoto *et al*, 2022). For example, the plant rhizosphere is inhabited by a wide range of bacteria, fungi, microalgae, viruses and archaea (Mendes *et al,* 2013), which protect the host against biotic and abiotic stresses, promote growth and increase nutrient availability (Chagas *et al,* 2018). Further, in humans and crops, microbiome dysbiosis is linked to severe chronic disease and stress (Peixoto *et al*, 2022). Key symbionts of cnidarian animals, such as reef-building corals and sea anemones, are dinoflagellate microalgae in the family Symbiodiniaceae (Davy *et al,* 2012). These symbionts translocate products of carbon fixation and nitrogen assimilation to the host (Burriesci *et al,* 2012; Kopp *et al,* 2015) and meet up to ~161% of the host’s basal metabolic energy requirements (Muscatine *et al,* 1984). In exchange, they gain protection and host-derived inorganic nutrients (Davy *et al*, 2012). Metabolites play a central role in host-microbe interactions by mediating and controlling the basic processes of recognition, signaling and communication (Song *et al,* 2015; Cleary *et al,* 2017; Chagas *et al,* 2018). In plants, for instance, lipids such as triacylglycerols, phospholipids, galactolipids, and sphingolipids are crucial for signaling and system-level energy storage (Bhattacharya, 2022)

To understand host-microbe interactions, tissue homogenates have traditionally been analyzed with liquid/gas chromatography coupled to mass spectrometry (LC/GC-MS) to measure the host/symbiont metabolite profiles. However, symbiotic microorganisms are often found within specific anatomical structures or cell types of the host. For instance, the sulfur-oxidizing and methane-oxidizing gammaproteobacterial symbionts of deep-sea mussels (*Bathymodiolus puteoserpentis*) from hydrothermal vents are only found in the organism’s epithelial cells (Geier *et al,* 2020). Similarly, the bioluminescent bacterium *Vibrio fischeri* is localized within the epithelial layers in the light organ of the Hawaiian bobtail squid (*Euprymna scolopes*) (Cleary *et al,* 2017). Spatial information is lost in tissue homogenates and the sensitivity for measuring specific foci of nutrient/metabolite exchange (e.g., within a particular body structure) are dampened in bulk analyses.

The loss of dimensionality substantially restricts the value of metabolomics to provide insights into the microbial ecology of symbioses, as it can limit biological inferences to be drawn from the data. For instance, Williams *et al.* (2021) utilized UHPLC-MS on tissue homogenates of the coral *Montipora capitata* in symbiosis with Symbiodiniaceae, and detected an enrichment of montiporic acids under elevated temperature. Montiporic acids are cytotoxic and antimicrobial compounds found in *M. capitata* eggs that are also known to reduce the photochemical efficiency of coral microalgal symbionts (Hagedorn *et al,* 2015). Without knowing whether the enriched montiporic acids were co-localized with the gonads or the gastrodermal cells that contain the microalgal symbionts, it is impossible to tell if this result is linked to coral sexual reproduction or symbiont photosynthesis. The lack of spatial information can also reduce the ability to detect significant differences between experimental treatments. Sphingolipids can accumulate in host microalgal symbiont-containing gastrodermal cells of the sea anemone *Exaiptasia diaphana.* However, contrary to expectations, LC-MS/MS analysis of homogenized tissues did not reveal a significant sphingolipid concentration difference between anemones exposed to different light/dark treatments, possibly due to the dilution effect of non-symbiotic cells (Kitchen *et al,* 2017). These examples highlight that spatial knowledge of molecular species is key for obtaining in-depth understanding of host-microbe interactions.

Nano-scale secondary ion mass spectrometry (NanoSIMS) is one technique to study host-microbe interactions spatially. NanoSIMS is an isotope and elemental imaging technique that, in combination with transmission electron microscopy, can visualize relative isotopic enrichment in biological samples at very high spatial resolution (up to 50 nm) (Nuñez *et al,* 2018; Siegel *et al,* 2018). Through ^13^C and ^15^N labelling, NanoSIMS has enabled researchers to explore question such as symbiont carbon assimilation and translocation (e.g., Ros *et al,* 2021), as well as symbiont nitrogen assimilation rate and nutrient competition *in situ* in corals (e.g., Krueger *et al,* 2020). However, the high energy monoatomic beam of NanoSIMS results in severe fragmentation of larger-sized molecules (e.g., proteins, lipids) and limits researchers to study very small molecules only (i.e., < 200 m/z); and the technique can only image seven molecules at once because of its mass analyser limit (Siegel *et al,* 2018). Further, NanoSIMS requires samples with a skeleton (e.g., scleractinian corals) to be decalcified prior to imaging (Krueger *et al*, 2020; Ros *et al*, 2021) and hence spatial information across the coral polyp is poorly preserved.

MSI metabolism, recently termed spatial metabolomics (Alexandrov, 2020), is one method to overcome the limitations of traditional metabolomics and NanoSIMS. With the exception of medical studies, the application of MSI in microbiology is in its infancy but its potential is vast. MSI utilizes intact tissue sections, allowing researchers to examine the spatial distribution of hundreds to thousands of molecules in complex biological samples *in situ* simultaneously. Unlike NanoSIMS, isotopic labelling or endoskeleton decalcification is not required for MSI. Through 2D rasterized ionization, the method generates thousands of charged species that are introduced into a mass spectrometer, capable of detecting a wide m/z range of 20 – 500k+. Each position of the 2D raster pattern collects all measurable mass-to-charge (m/z) values within a fixed range, which relate to molecular species. These positional spectra (termed pixels) are summed across the whole 2D area, where ion density maps can be generated to visualize the spatial location and relative intensity of each metabolite (Siegel *et al,* 2018). While the metabolite peak area itself does not reflect absolute concentrations (as this depends on e.g., ionization efficiency, transmission efficiency through the mass spec, etc.), the peak area scales linearly with metabolite concentration and comparison of relative intensity between samples can be made (Liu & Locasale, 2017). Further, absolute quantitation can be achieved through the use of external/internal standards (Unsihuay *et al,* 2021). The spatial information provided by MSI has been proven highly valuable in medical studies, particularly for drug discovery, disposition and disease state assessment (Roberts *et al,* 2022; Siegel *et al,* 2018). However, MSI is underutilized beyond the medical research field.

The ability to study hundreds to thousands of metabolites *in situ* via MSI makes this a powerful tool for researchers to decipher host-microbe interactions. Only a few studies have explored the value of MSI to examine host-bacteria interactions. Examples are the mapping of the spatial metabolome of a deep-sea mussel which has intracellular bacteria embedded within epithelial cells (Geier *et al*, 2020); investigating the role of bacteria-derived secondary metabolites in larval maturation of a marine snail (Rudd *et al,* 2015); and bacteria-derived prophylaxis in wasp cocoons (Kroiss *et al*, 2010). MSI has provided invaluable insights and new discoveries in these studies, such as the discovery of specialized metabolites at the host–microbe interface specific for mussel-methane-oxidizing bacterial interactions (Geier *et al*, 2020). Equally, MSI has the potential for elucidating the cnidarian-dinoflagellate symbiosis - an avenue yet to be explored. Coral bleaching is the loss of microalgal symbionts (Symbiodiniaceae) that often results in host mortality (Hughes *et al*, 2017, 2018). With < 2% of the Great Barrier Reef (GBR) having escaped bleaching since 1998 (Hughes *et al,* 2021), knowledge on host-symbiont interactions is critical to develop novel reef conservation and restoration interventions.

Obtaining intact tissue sections to investigate cnidarian-Symbiodiniaceae interactions *in situ* with MSI is challenging. Typical cryosectioning techniques cannot maintain the natural shape of delicate soft-bodied cnidarians (e.g., sea anemones); nor are these methods able to cut through the CaCO_3_ skeleton of hard-bodied cnidarians (e.g., scleractinian corals) without disrupting its tissue. We have overcome these challenges and present a spatial metabolomics workflow that can reveal host and Symbiodiniaceae metabolite profiles *in situ* at 50 μm resolution using intact tissue sections of a soft-bodied (the sea anemone, *E. diaphana*) and hard-bodied cnidarian (the scleractinian coral, *Galaxea fascicularis).* To verify the biological application of the method, we test 1) if the metabolite distribution is linked to specific host anatomical structures and 2) if symbiont identity drives anemone host metabolism in a spatial manner. To examine the effects of symbiont identity on the host, *E. diaphana* of the same genotype was rendered aposymbiotic (i.e., free of microalgal symbionts) with menthol bleaching and inoculated with the homologous *Breviolum minutum* microalgal symbionts (hereafter referred to as B1-anemones), or the heterologous *Cladocopium* C1^acro^ microalgal symbionts (hereafter referred to as C1^acro^-anemones) (Tsang Min Ching *et al*, 2022). These *E. diaphana* have been confirmed to be in symbiosis with *B. minutum* and C1^acro^, respectively, using ITS2 metabarcoding (Tsang Min Ching *et al,* 2022).

## RESULTS AND DISCUSSION

### A spatial metabolomic workflow for marine invertebrates

Spatial omics studies are rare in marine invertebrates (Geier *et al,* 2020; Goto-Inoue *et al,* 2020; Hamilton *et al,* 2022), and the difficulty in obtaining intact tissue sections in the organism’s natural shape is a major hurdle. Through the combination of anesthetizing, embedding and cryofilm application, the cryosectioning workflow presented produced intact tissue sections for a soft-bodied (sea anemone *E. diaphana*) and a hard-bodied (coral *G. fascicularis*) cnidarian in their natural shape (Fig. 1); this is a prerequisite for spatial metabolomics and other spatial analysis. This method is applicable to other soft- and hard-bodied marine invertebrates, opening up new opportunities for the broader marine invertebrate research field. All detected metabolites are immediately contextual, in that they can be correlated to host/symbiont anatomical structures with known ecological functions. Compared to NanoSIMS, which is restricted to imaging seven ions at a time, a total of 631 metabolites (after background signal removal) were imaged in this study across the anemone–symbiont metabolome using MALSI-MSI (Supplementary dataset S1). Of these, 208 (33%) were annotated, falling within 31 major groups covering structural, energetic and signaling metabolites (Supplementary dataset S1, Table S1). The remaining 67% represents ambiguous or unknown metabolites, alluding to a considerable ‘dark metabolome’ in cnidarian tissues. Annotation in this space is still one of the major bottlenecks in MSI experiments, but this study has provided the first step to fill this knowledge gap.

**Fig. 1.**
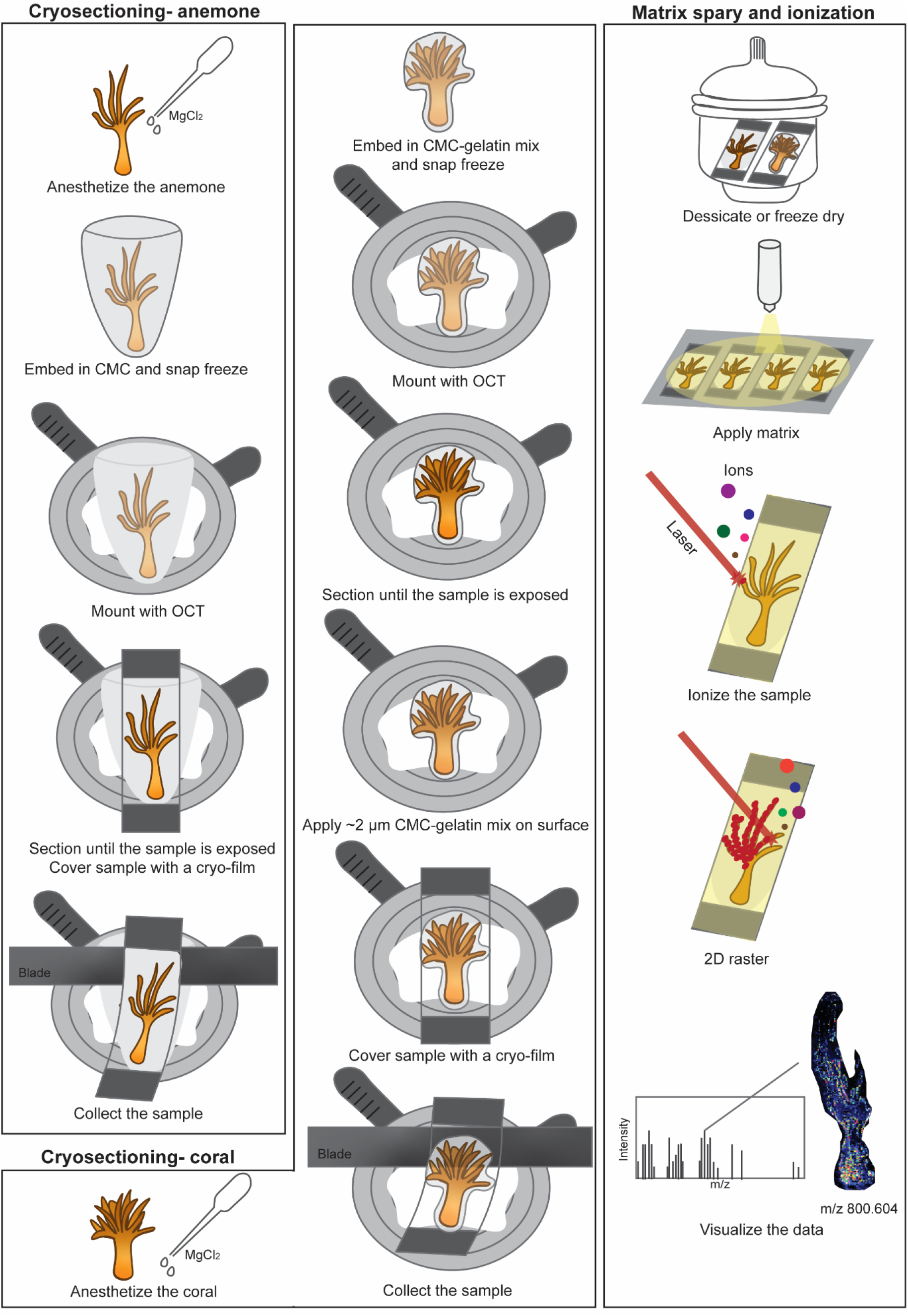
A spatial metabolomics method. Cryosectioning and matrix-assisted laser desorption/ionization mass spectrometry imaging (MALDI-MSI) workflow for the soft-bodied sea anemone *E. diaphana* and the hard-bodied coral *G. fascicularis.* Abbreviations refer to: CMC: 2% carboxymethyl cellulose, OCT: optimal cutting temperature compound.

### MALDI-MSI reveals spatial distribution patterns of metabolites that link to functionality

#### Tissues surrounding the gastrovascular cavity as a focal site of host-symbiont regulation

MSI revealed the localization of metabolites that would otherwise be lost in tissue homogenates. Among the 31 metabolite groups detected in anemones, ceramides (group Cer, GlcCer, HexCer; 26 of the major chemical species detected) were found to be spatially concentrated in the tissues surrounding the gastrovascular cavity and occasionally the column outer wall and actinopharynx of symbiotic anemones (Figs. 2a, b, c). In contrast, betaine lipids (group DGCC, MGCC, DGTS; 13 of the major chemical species detected) were primarily concentrated in the tentacles of symbiotic anemones (Figs. 2a, b, c). Interestingly, the ratio between these metabolite groups showed a clear pattern that separates anemones based on the identity of their Symbiodiniaceae symbiont (Fig. 2c), indicating that symbiont identity can change system-level host metabolism (see section “Symbiont identity drives system-level host metabolism spatially”).

**Fig. 2.**
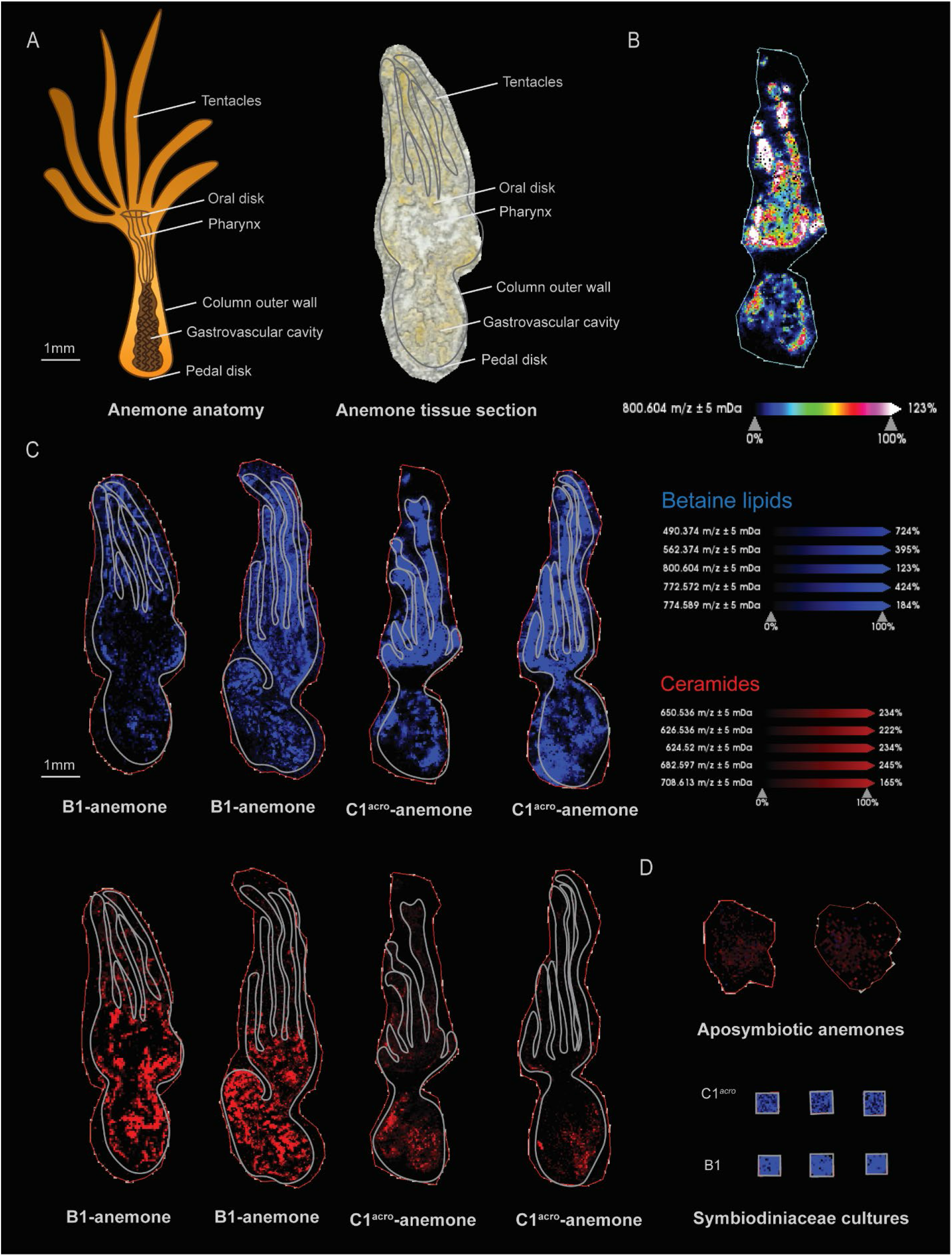
Spatial metabolite distribution pattern in sea anemones. (**A**) Anatomy and a tissue section of the sea anemone *E. diaphana.* (**B**) Spatial intensity of the betaine lipid DGCC m/z 800.604 with a coloured metabolite intensity scale, as an example of the intensity pattern relevant to all other betaine lipids. (**C**) Spatial distribution of five selected betaines lipids (blue, concentrated in the tentacles) versus five selected ceramides (red, concentrated within the tissues surrounding the gastrovascular cavity of symbiotic anemones) in anemones in symbiosis with *B. minutum* (B1-anemones) and C1^acro^ (C1^acro^-anemone). (**D**) Betaines lipids (blue) and ceramides (red) relative intensities in the Symbiodiniaceae cultures and aposymbiotic anemones. Note that aposymbiotic anemones are small compared to symbiotic ones since the absence of microalgal symbionts resulted in reduced nutrient supply and hence anemone growth.

The concentration of ceramides within the tissues surrounding the gastrovascular cavity reflects it is the primary site of regulation of Symbiodiniaceae symbionts by the cnidarian host. Ceramides are intermediates in the biosynthesis and metabolism of all sphingolipids — a class of lipids that are important constituents of biological membranes which function in cell signaling through the activation of specific G-protein-coupled receptors — regulating processes such as apoptosis, cell survival, inflammation, autophagy and oxidative stress responses (Bhattacharya, 2022; Rosset *et al*, 2021). The sphingosine rheostat refers to the balance in cellular concentrations of pro-survival sphingolipids (i.e., sphingosine-1-phosphate; S1P) and pro-apoptotic sphingolipids (i.e., sphingosine and ceramide), which is a homeostatic process that determines cell fate (Kitchen *et al*, 2017; Rosset *et al,* 2021) (Fig. 3). Pro-survival and pro-apoptotic sphingolipids are interconvertible by the catalytic activities of sphingosine kinase (SPHK) and S1P phosphatase (SGPP); where SPHK promotes the conversion to S1P and enhances cell survival and proliferation, and SGPP promotes the conversion to sphingosine that activates pro-apoptotic cellular cascades (Fig. 3). The regulatory mechanisms that balance sphingosine conversion could be features that determine resilience to and recovery from abiotic stress such as coral bleaching caused by ocean warming, which is a global concern for reef-building corals.

**Fig. 3.**
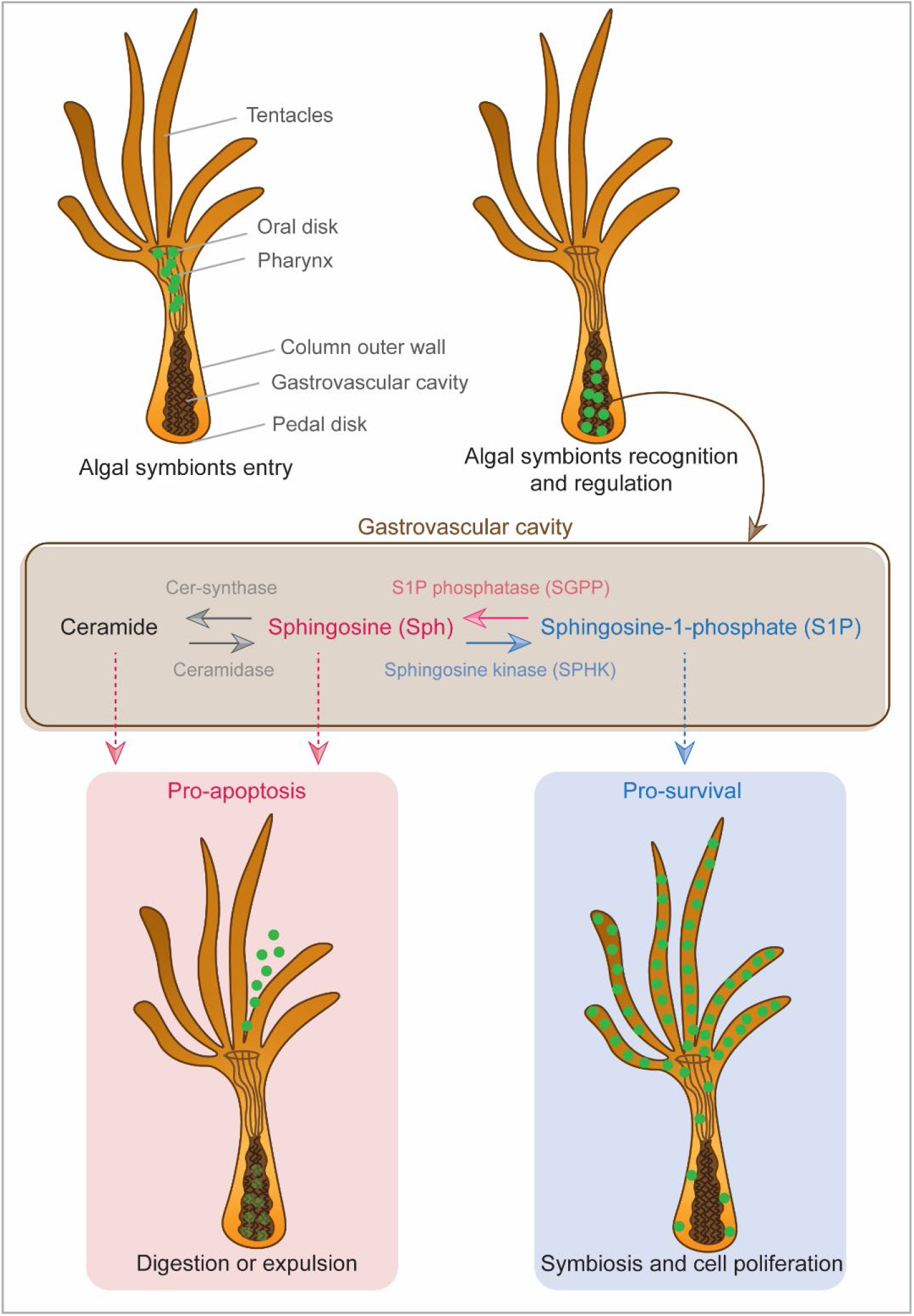
The cnidarian sphingosine rheostat. The mode of entry of microalgal symbionts into a cnidarian host and the potential regulatory role of sphingolipids in the cnidarian-Symbiodiniaceae symbiosis.

While several previous studies have pointed towards a regulatory role of sphingolipids in cnidarian-Symbiodiniaceae symbiosis, none has explored their spatial distribution and associated implications. Anemones supplied with exogenous pro-survival sphingolipids (S1P) exhibited less bleaching under elevated temperatures compared to control anemones (Detournay & Weis, 2011), and the pro-apoptotic SGPP gene is downregulated in symbiotic compared to aposymbiotic anemones (Kitchen *et al,* 2017; Rodriguez-Lanetty *et al,* 2006). Uptake of Symbiodiniaceae symbionts by cnidarian hosts occurs in the gastrovascular cavity, where they are incorporated into the host’s gastrodermal cells via phagocytosis (Davy *et al,* 2012). A high concentration of ceramides in the tissues within the gastrovascular cavity supports the role of sphingolipids in cnidarian-Symbiodiniaceae symbiosis regulation, and indicates that this is likely where the fate of the microalgal symbiont (i.e., being digested/expelled or incorporated as a symbiont) is determined by the sphingosine rheostat (Fig. 3). Further, how the cnidarian sphingosine rheostat may be affected by the microalgal symbiont species or strain harboured has previously not been explored.

This study demonstrates that the relative intensity of the majority of ceramides are significantly higher in B1-anemones (with homologous symbionts) than C1^acro^-anemones (with heterologous symbionts) (Figs. 2c, S1, Table S2); indicating that symbiont identity can drive spatial changes in host ceramide level that governs the sphingosine rheostat. Alternatively, this difference could be driven by the cell density and growth rates of the microalgal symbionts. At six months post microalgal symbiont inoculation to the anemones, B1-anemones had higher cell density than C1^acro^-anemones (referred to as “WT10” in (Tsang Min Ching *et al*, 2022)), and their cell densities were eventually equal by ~1.5 year post inoculation) (Tsang Min Ching *et al,* 2022). The samples of this study were collected at 10 months post inoculation, therefore the symbiont cell density was likely higher in B1-anemones than C1^acro^-anemones. Hight symbiont cell density and/or faster symbiont growth rate may require more cells to be removed to maintain a stable and healthy symbiont density in the host, which may have been facilitated by higher ceramide relative intensity.

#### Tentacles as the primary site for photosynthesis

The distribution pattern of betaine lipids suggests that once acquired, microalgal symbionts are mostly transported to the light-exposed anemone tentacles for photosynthate production (Figs. 2b, c). We also investigated the spatial distribution of these betaine lipids in symbiotic corals and found the same pattern, indicating that this is likely a shared character among cnidarians (Fig. 4). Betaine lipids are absent in higher plants, but are known to occur in a wide range of marine microalgae (including Symbiodiniaceae (Kato *et al,* 1996; Roach *et al,* 2021)), with the three main microalgal betaine lipids being DGTS, diacylglyceryl hydroxymethyl-N,N,N-trimethyl-beta-alanine (DGTA) and diacylglyceryl carboxyhydroxymethylcholine (DGCC) (Cañavate *et al,* 2016; Kato *et al,* 1996). Betaine lipids were abundant in the Symbiodiniaceae cultures and in symbiotic anemones, but were absent in aposymbiotic anemones (Figs. 2c, d, Table S3). The only diacylgyceryl-N-trimethylhomoserine (DGTS) betaine lipid detected (m/z 472.363) occurred in high relative intensity in all Symbiodiniaceae cultures but was nearly absent in symbiotic anemones. This suggests that the metabolic state of the microalgal symbionts underwent changes when shifting from the free-living to the symbiotic stage. Conversely, this betaine lipid had high relative intensity in symbiotic corals (Fig. 4). Since the microalgal symbiont identity differed between the anemones (*Cladocopium* C1^acro^ or *Breviolum*) and coral (*Cladocopium* C40), the data reflect that these symbionts may have different lipid constitution or behave differently to each other in the free-living versus symbiotic life stage.

**Fig. 4.**
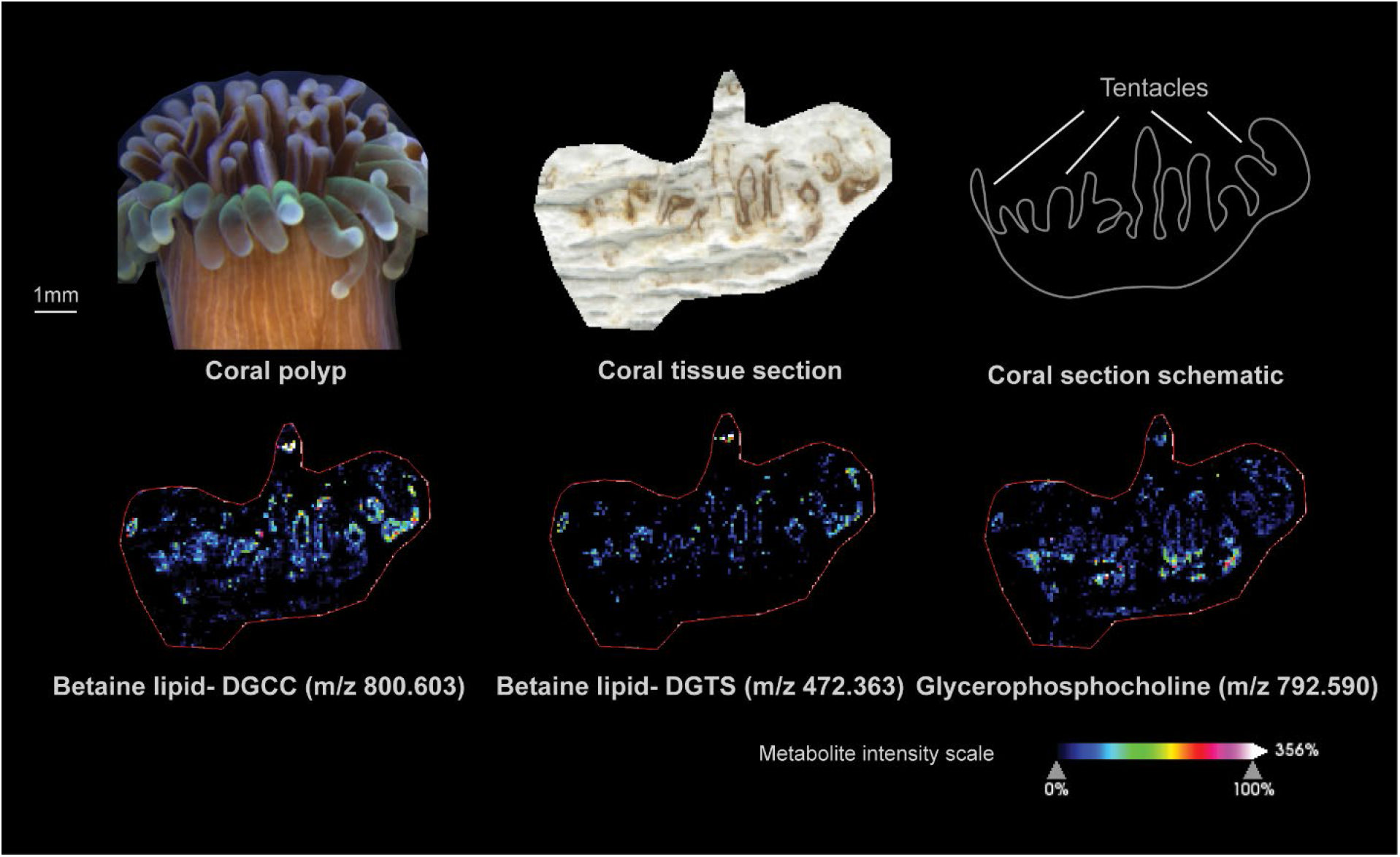
Spatial distribution of three selected metabolite in a single polyp of the coral *G. fascicularis*. Note the concentration of betaine lipid (microalgal signal) throughout the tentacles of the symbiotic coral and concentration of glycerophosphocholine (PC, host signal) at the base of the coral tentacles.

In the sea anemones *Condylactis gigantea* and *Anthopleura elegantissima,* the majority of Symbiodiniaceae occur in the tentacles, or tentacles and oral disk (Augustine & Muller-Parker, 1998; Dykens & Shick, 1984; Kellogg & Patton, 1983). Consistent with this notion, Dykens & Shick (Dykens & Shick, 1984) observed much higher chlorophyll content and oxygen production in the anemone’s tentacles and oral disk than its body column and pedal disc. In *E. diaphana,* while no direct observation of Symbiodiniaceae concentration in the tentacles has been previously reported, Symbiodiniaceae colonization in aposymbiotic anemones starts in their oral disk and tentacles, followed by the body column and finally the pedal disk, regardless of the taxonomic identity of the Symbiodiniaceae (Gabay *et al,* 2018). The higher relative intensity of betaine lipids in the tentacles of *E. diaphana* compared to the body observed here is in line with the spatial distribution pattern of Symbiodiniaceae symbionts in other anemone species. This spatial distribution may reflect preference by the host or microalgae due to the varying *in hospite* light environments, where greater exposure to light in the tentacles is ideal for photosynthesis.

A recent single-cell RNA sequencing study showed unique gene expression patterns in different coral cell types (Levy *et al*, 2021). Fluorescence-activated cell sorting in the coral *Stylophora pistillata* was able to separate Symbiodiniaceae-hosting gastrodermal cells from non-symbiotic gastrodermal cells; single-cell RNA sequencing revealed Symbiodiniaceae-hosting gastrodermal cells were enriched in 353 host genes including genes related to lipid metabolism, carbonic anhydrases, amino acid and peptide transporters. In addition, only Symbiodiniaceae-hosting gastrodermal cells expressed all the mRNAs for the galactose-catabolism Leloir pathway, indicating the translocation of galactose from the symbionts to the coral host was occurring within this cell type. As in the present study, these findings allow researchers to link enrichment of a gene product or metabolite to specific cell compartments or structures and make biological inferences on their functionality, highlighting the value of spatial metabolomic and spatial gene expression analysis. The spatial metabolite pattern revealed in MSI studies can also be used to select areas-of-interest to further explore specific research questions. For instance, anemone researchers interested in symbiont regulation and sphingosine rheostat of the host can selectively sample the tissues surrounding the animal’s gastrovascular cavity, whereas researchers interested in photo-physiology and oxidative stress of the microalgal symbionts can target the animal’s tentacles, where focal changes are most likely to occur in the metabolome and the expression of genes.

### Sphingosine rheostat was likely inactive in aposymbiotic anemones

Aposymbiotic anemones were in a different metabolite state compared to symbiotic anemones, with 249 out of 631 metabolites (39.5%) showing statistically significant relative intensity (Figs. 2c, d, 5, Table S4). Compared to symbiotic anemones and regardless of their Symbiodiniaceae symbionts’ identity, aposymbiotic anemones lacked betaine lipids and had a low relative intensity of ceramides (CER) and diglycerides (DG) in the host tissues (Fig. 5d). However, aposymbiotic anemones had a much higher relative intensity of many phosphatidylcholine (PC) lipids in the host component, which dominated the aposymbiotic anemone metabolome (Fig. 5d). The Symbiodiniaceae cultures were rich in the 13 major detected chemical species of betaine lipid, but had no or minimal relative intensity in ceramides (Fig. 2d). The absence of betaine lipids in aposymbiotic anemones (Fig. S2, Table S3) confirms that these lipids are produced by the microalgal symbionts and that the signals detected in the anemone tissues were of microalgal symbiont origin. The low relative intensity in ceramides in aposymbiotic anemones and Symbiodiniaceae cultures (Fig. 2d) suggests the high ceramide intensities detected in symbiotic anemones is a consequence of symbiosis, where the presence of symbionts may have triggered *de novo* synthesis or salvage of sphingolipids in the host that activate the cnidarian-Symbiodiniaceae regulatory system. Alternatively, this regulatory system could be based on a shared metabolic pathway, where the host and symbiont each have parts of the pathway and could only be activated when both parties are present.

**Fig. 5.**
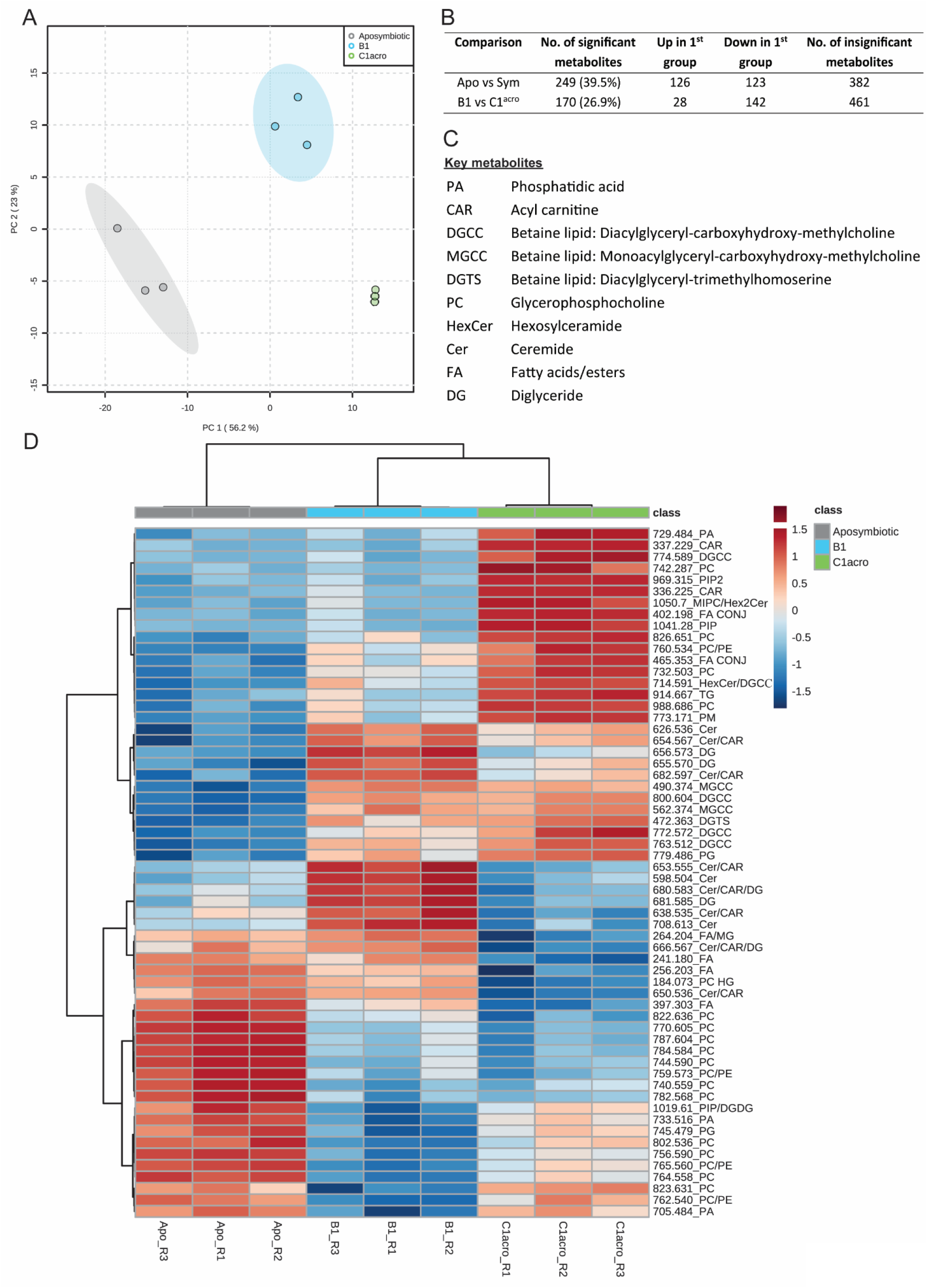
Statistical comparisons of aposymbiotic, B1- and C1^acro^ anemones. (**A**) PCA using all 631 metabolites. Note that the 95% confidence ellipse for the three C1^acro^ samples (green) is not visible at the scale presented due to the close proximity of the data points. (**B**) Pairwise comparison results using all 631 metabolites. A metabolite is considered significant when FDR < 0.05 and FC > 2.0. (**C**) Abbreviation and full names of key metabolites. (**D**) Heatmap of the top 60 significantly different metabolites in an ANOVA between aposymbiotic anemones, anemones in symbiosis with the homologous B1 (*B. minutum*) and anemones in symbiosis with the heterologous C1^acro^. Only metabolites with an annotation are shown here. See supplementary Table S1 for full names of all metabolite abbreviations. “R” in the sample name refers the replicate number.

### Symbiont identity drives system-level host metabolism spatially

The metabolome of C1^acro^-anemones was distinct from that of B1-anmeones, with 170 (26.9%) significantly different metabolites (Figs. 2c, 5, Table S5). Compared to B1-anemones, C1^acro^-anemones had lower relative intensity in many ceramides and fatty acids, as well as several diglycerides (DG), phosphatidylcholines (PC) in the host component (Fig. 5d). Of the 26 major ceramides-related chemical species detected, C1^acro^-anemones (with heterologous symbionts) had a lower relative intensity in almost all ceramides, but higher relative intensity in nearly all the higher molecular weight hexosylceramide (HexCer) and mannosylinositol phosphorylceramide (MIPC) (Fig. S1, Table S2). Of the 13 major chemical species detected for betaine lipids, C1^acro^-anemones had significantly higher relative intensity in two metabolites (m/z 772.572, 774.589, match to polyunsaturated structures) compared to B1-anemones (Fig. S2, Tables S3, S6). C1^acro^-anemones had lower relative intensity in some fatty acids in the host component (Fig. 5d). A combined transcriptomic and metabolomic study has demonstrated that *E. diaphana* associated with heterologous microalgal symbionts showed an up-regulation of host innate immunity metabolites and genes, increased lipid catabolism and decreased transport of fatty acids to the host, whereas those associated with homologous microalgal symbionts showed immunotolerance and symbiont-derived nutrition (Matthews *et al*, 2017). The lower host relative intensity in fatty acids in C1^acro^-anemone (with heterologous symbionts) compared to B1-anemones (with homologous symbionts) is consistent with the above study and in agreement with a previous observation on the same anemones that heterologous symbionts can be slightly less nutritionally beneficial to their host (Tsang Min Ching *et al*, 2022).

In summary, we have demonstrated a MALDI-MSI workflow that can map *in situ* host and symbiont metabolites with relevance to host anatomic structures in soft- and hard-bodied cnidarians. The approach allows researchers to make biological inferences about functionality that would have been overlooked with traditional metabolomics based on bulk tissue analysis, and the methodology is applicable to other marine invertebrates with a delicate soft body or with a hard skeleton, as well as to other symbiotic systems. This is the first study to reveal the cnidarian-Symbiodiniaceae metabolome *in situ.* Using the cnidarian-Symbiodiniaceae symbiosis model of the sea anemone *E. diaphana,* MALDI-MSI demonstrated how microalgal symbiont identity can drive system-level change in host metabolism spatially. Questions around the role of a symbiont on host fitness within a given environment are best addressed by sensitive measurements at the specific foci of nutrient exchange. Areas-of-interest identified in MSI can be isolated (e.g., using laser microdissection) for detailed characterization, such as metabarcoding to profile the microbial community and metagenomics to examine functional potential (Wada *et al*, 2022). With the increasing amount of evidence of a shift in host-microbe dynamics from mutualism to commensalism to parasitism/pathogenicity under a changing climate across ecosystems (Baker *et al*, 2018; Chagas *et al*, 2018), novel insights into host-microbe interactions provided by spatial metabolomics is becoming increasing important.

## MATERIALS AND METHODS

### Experimental design and sample collection

For the soft-bodied cnidarians, GBR-sourced *Exaiptasia diaphana* (genotype AIMS4) in symbiosis with the homologous *Breviolum minutum* (B1-anemones, which were inoculated with symbiont culture SCF 127-01, ITS2 profile: B1-B1o-B1p), or the heterologous *Cladocopium* C1^acro^ (C1^acro^-anemones, which were inoculated with symbiont culture SCF 055-01.10, ITS2 profile: C1-C1b-C1c-C42.2-C1br-C1bh-C1cb-C72k) were used (Table S7, Fig. S3). Their symbiont identity was confirmed by ITS2 metabarcoding six months prior to sampling for MSI (Tsang Min Ching *et al*, 2022) and reverified three months post-sampling that there was no change (Sakamoto, 2021). *E. diaphana* were sampled 10 months post-inoculation for MSI and were maintained under ambient temperature of 27 *°C* at 30 μmol m^-2^ s^-1^ (12:12h, light: dark). For hard-bodied cnidarians, the coral *G. fascicularis* in symbiosis with *Cladocopium* C40 (ITS2 profile: C40-C3-C115-C40h) was obtained from Palm Islands (Fig. S3), GBR, and kept under ambient temperature of 27 °C and at 130-150 μmol m^-2^ s^-1^ at full sun (12:12h light: dark). *E. diaphana* and *G. fascicularis* were fed five days a week with freshly hatched *Artemia nauplii* or microalgae mix respectively. To avoid metabolite contamination from the food source, feeding was stopped five days prior to sampling. Three samples were collected for each cnidarian group (i.e., B1-anemone, C1^acro^-anemone, aposymbiotic anemone and *G. fascicularis*) Aposymbiotic (i.e., microalgal symbiont free) anemones were produced by a modified menthol bleaching method detailed in Tsang Min Ching *et al.* (2022, modified from Matthews *et al*, 2016). The steps for sample collection were:

1. For *E. diaphana,* singles polyps were collected using a disposable pipette.
2. For *G. fascicularis*, single coral polyps were removed with a bone cutter and left to rest in the aquaria for seven hours before fixation (Table 1).
3. The cnidarians were each placed in an individual well of a 12-well plate with 2 mL of seawater and left in the dark for 30 min to allow them to relax and fully extend their tentacles.
4. To avoid mucus formation during the sampling process (which would hinder MS ionization), cnidarians were anaesthetized with 1 mL of 0.4 M MgCl_2_ and left in the dark for a further 30 min.
5. Anaesthetized cnidarians were then rinsed twice in MQ water and carefully dried with Kimwipes and embedded to maintain the natural shape of the animals for cryo-sectioning.
6. For *E. diaphana*, each sample was placed in an embedding container with 1 mL of embedding media (2% carboxymethyl cellulose, CMC) and snap frozen on dry ice (Fig. 1).
7. The CaCO_3_ skeleton of *G. fascicularis* cryo-sectioning required optimization, hence a pilot study was conducted with a range of embedding media (CMC, agar, CMC mixed with gelatin, CMC mixed with agar) at different concentrations to identify the most suitable embedding media. The media that consistently yielded the most intact coral sections (i.e., a ratio of 10 mL 2% CMC: 0.123 g of gelatin) was then used for embedding of the samples to be subjected to MSI. The *G. fascicularis* polyps were dipped into the CMC-gelatin mix with a pair of forceps and snap frozen on dry ice.
8. Cnidarian samples were wrapped in aluminum foil and stored at −80 °C until further processing.

**Table 1.** Sample preparation and MALDI-MSI parameters of the sea anemone *E. diaphana* and the coral *G. fascicularis.*

|  | Sea anemone *E. diaphana* | Coral *G. fascicularis* |
| --- | --- | --- |
| **Sampling** | Collect directly with a disposable pipette | Remove with a bone cutter and leave to rest |
| **Embedding media** | 2% CMC | 2% CMC with gelatin (10 mL CMC: 0.123 g gelatin) |
| **Embedding method** | Place in an embedding container with 1 ml CMC | Dip sample into the CMC-gelatin mix with a pair of forceps |
| **Cryosectioning °C** | -20 °C | -25 °C |
| **Cryosectioning blade** | Low-profile stainless-steel disposable blade | High profile disposable diamond blade |
| **Desiccation method** | Leave in desiccator for 30 min | Freeze dry for 12 min |
| **Matric coating** | 100 µL min<sup>-1</sup> for four passes | 70 µL min<sup>-1</sup> for six passes |
| **Laser power** | 38% | 52% |
| **Laser shot** | 250 | 500 |

### Cryosectioning and matrix spray

Cryosectioning was conducted with a Leica CM 1860 at −20 °C (*E. diaphana*) or −25 °C (*G. fascicularis*) (Fig. 1, Table 1) with the following steps:

1. Samples were mounted on the cryostat with an optimal cutting temperature compound (OCT), while ensuring that the tissue sections for imaging were not contaminated by OCT.
2. Samples were sectioned at 12 μm thickness until arriving to the middle of the animal, where maximum areas of the frozen body and tentacles were exposed. A low-profile stainless-steel disposable blade was used for the soft-bodied *E. diaphana* and a more robust high profile disposable diamond blade was employed for the hard-bodied *G. fascicularis*.
3. A cryofilm was then attached to the exposed sample and a tissue section was collected at 12 μm thickness onto the cryofilm (Fig. 1).
4. To minimize scratches on the tissue sections caused by scarping of the CaCO_3_ skeleton during sectioning in *G. fascicularis,* a thin layer (~2 μm) of the CMC-gelatin mix was applied on the exposed sample surface to strengthen its integrity before collecting a section.
5. Four consecutive sections were collected per sample.
6. For *E. diaphana*, tissue sections were placed in a desiccator for 30 min under a small amount of dry ice, which allowed the sections to gradually come to room temperature during desiccation.
7. For *G. fascicularis,* the desiccator method was insufficient to maintain tissue integrity, and the tissue sections were instead freeze dried for 12 min under ~-50 °C and ~0.08 mbar.

Matrix application was carried out with the following steps:

1. Two desiccated sections per sample were mounted on a stainless-steel sheet with carbon tapes and coated with a matrix (A-Cyano-4-hydroxycinnamic acid, HCCA) to assist ionization (Fig. 1).
2. For each matrix spray run, 30 mg of HCCA was dissolved in 6 mL of solvent (70% Acetonitrile, 30% H_2_O with 0.1% Trifluoroacetic acid) and loaded into the HTX TM-Sprayer™.
3. Matrix coating was conducted under 75°C at the flow rate of 100 μL min^-1^ for four passes (*E. diaphana),* or 70 μL min^-1^ for six passes (*G. fascicularis).*
4. Tissue sections were desiccated in a desiccator for a further 20 min before imaging (Table 1).

### MALDI-MSI analysis

MALDI-MSI analysis was performed on a Bruker SolariX (7T XR hybrid ESI-MALDI-FT-ICR-MS) with a mass resolving power of 200000 and equipped with a SmartBeam II UV laser. Prior to data collection, the instrument was calibrated with a red phosphorus standard to ensure that its mass error was less than 1.5 ppm. Two technical replicates were imaged per sample and two samples were imaged per MSI run. The area of interest for imaging was defined using flexImaging 4.1 (Bruker Daltonics) and data acquisition was controlled via Bruker Daltonics ftmsControl 2.1.0. Spectra were collected at a spatial resolution of 50 μm and a range of 150-2000 m/z under positive ion mode. Laser diameter and power were set to 45 μm and 38% (*E. diaphana*) or 52% (*G. fascicularis);* and a total of 250 (*E. diaphana*) or 500 (*G. fascicularis*) laser shots were applied at each 50 μm pixel at a frequency of 2 kHz.

### Data processing

Data processing involved:

1. Raw spectra files (.mis) were uploaded to SCiLS and combined into a single file (.slx) with the intervals set to ± 0.5 mDa (Supplementary dataset S2-S5).
2. A m/z peak list was generated using the sliding window function and a threshold cutoff of 9000 was applied, where the majority of the noises were removed.
3. To remove background peaks (contributed by e.g., instrument noise, CMC or gelatin instead of the biological sample), a region-of-interest (ROI) was created on the background area to identify and remove those associated peaks. A total of 631 peaks remained afterward denoising and each was visualized on SCiLS to confirm that they were associated with the biological samples (Supplementary dataset S1).
4. The intensities (average peak area) of the 631 peaks of each technical replicate were exported as excel files, using root mean square to normalize potential intensity differences between MSI runs.

Since each tissue section varied slightly in 2D area, the intensity values were normalized to the total surface area of the section to account for size differences:

1. The surface area of a tissue section was identified in SCiLS by overlaying the ion density maps of the most prominent peaks associated with the anemone host (m/z 792.590, glycerophosphocholine) and Symbiodiniaceae (m/z 800.604, betaine lipid DGCC) (Figs. 2c, 4). The use of these biological peaks accurately identified the tissue area, allowing gaps within a section (e.g., the empty space between 2 anemone tentacles) to be excluded from the surface area calculation (Fig. S4).
2. The ion density maps were exported to ImageJ and converted to 8-bit, and the ‘biological’ surface area was calculated by applying a signal threshold (consistently applied at 30).
3. The intensity of each peak was divided by the surface area of the section for normalization.
4. Technical replicates were then combined and the mean intensity of a peak was used for statistical analyses.

### Metabolite annotation

The peak list (631 peaks) was exported from SCiLS and peaks were initially annotated with LipidMaps, ChEBI and HMDB databases within METASPACE, with reference known coral-symbiont metabolites (Roach *et al*, 2021). Peak annotations were then verified using MALDI-thin layer chromatography (MALDI-TLC) to add a retardation factor (Rf), in lieu of retention time, and maintain the same ionization mode for analysis (laser desorption/ionization) (Figs. 6a, b). Select metabolites from Rf spots were analyzed using MALDI-MS/MS, laser induced dissociation (LID), to match to LC-MS/MS fragmentation patterns for identification (Fig. 6c). For TLC analysis, 1 mL of 1 x 10^6^ cells of the two Symbiodiniaceae cultures and two aposymbiotic anemones were included. Symbiodiniaceae cultures were collected and centrifuged at 1000 rcf for 5 min to remove the culture media. The pellets were resuspended in 1 mL MQ water and centrifuged again at 1000 rcf for 5 min. The supernatant was removed and the pellets were snap frozen on dry ice. Aposymbiotic anemones were rinsed in MQ water and dried with Kimwipes, before being snap frozen on dry ice. For the extraction, 500 μL of extraction buffer (CHCl_3_: MeOH, 2: 1) was added to the frozen samples, which were then sonicated and vortexed for 20 min. Fifty μL of LCMS grade water was added to the sample and centrifuged at 1000 rcf for 5 min. A total of 200 μL of a sample (the bottom phase) was pipetted to a new Eppendorf tube and dried under N2 flow. Samples were resuspended in 5 μL of CHCl_3_ and 1 μL of which was spotted on a TLC plate. The plate was allowed to develop for 30 min in a glass beaker with a mobile phase solvent consisting of 1.4 μL of LC-MS grade CHCl_3_, EtOH, H_2_O and 0.28 μL of triethanolamine (TEA). The plate was desiccated for 30 min, before being sprayed with HCCA and analyzed with MALDI-MSI with the aforementioned methods (with the exception of spatial resolution, laser diameter and power, which were set to 200 μm, 90 μm and 60% respectively). MALDI-LID-MS/MS was conducted on a Bruker Ultraflextreme MALDI-TOF, with a tuned LIFT method based on the lower mass range.

**Fig. 6.**
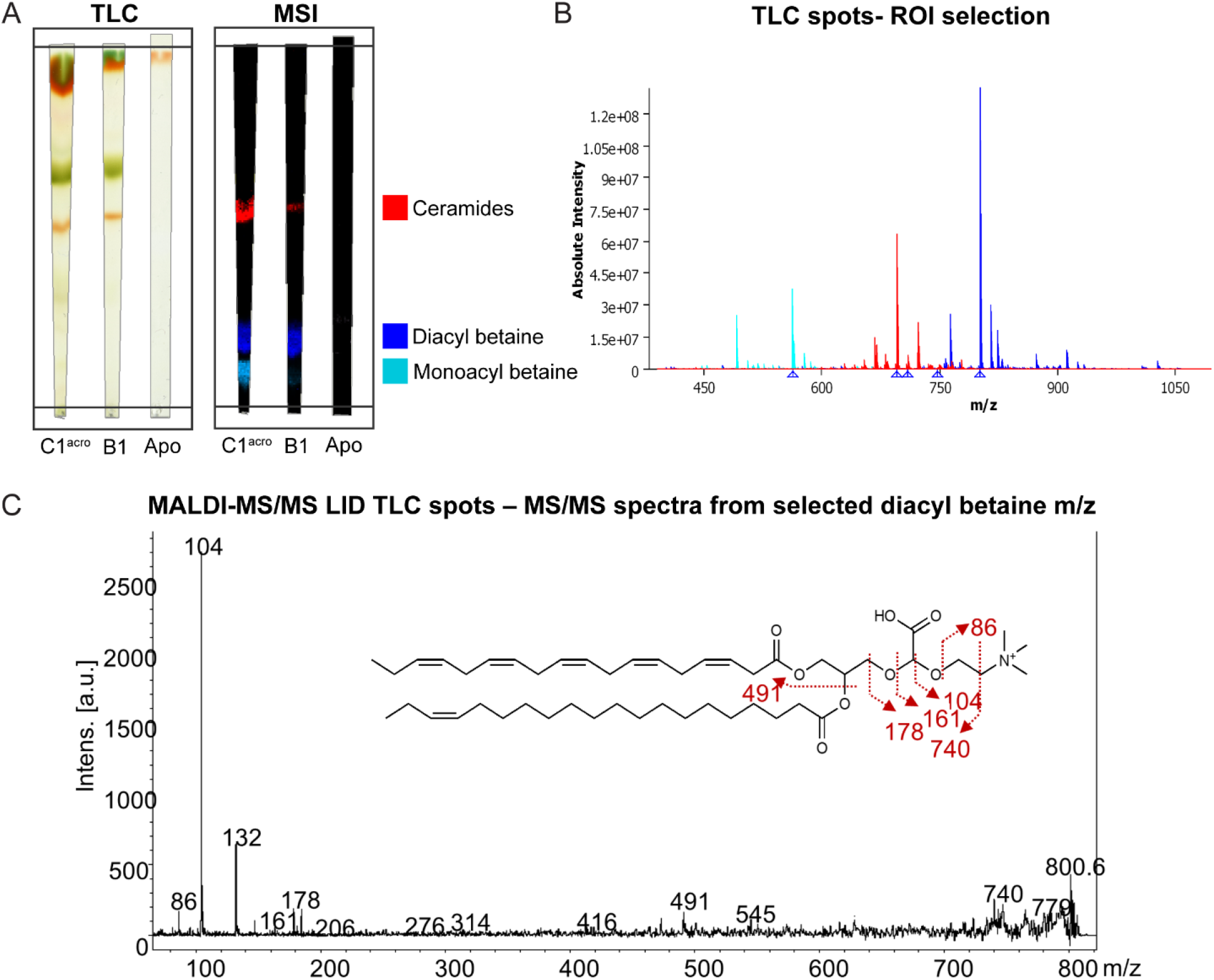
Example workflow for metabolite annotation. (**A**) Thin layer chromatography (TLC) and mass spectrometry imaging (MSI) for metabolite annotation on the algal culture *Cladocopium* C1^acro^ (C1^acro^) and *Breviolum minutum* (B1), as well as aposymbiotic anemones (Apo). (**B**) Spectra of three selected metabolite groups (i.e., ceramides, diacyl betaine lipids, monoacyl betaine lipids) based on region-of-interest (ROI) selection on the TLC. (**C**) Example MS/MS spectra of a diacyl betaine lipid generated using laser induced dissociation (LID).

### Statistical analysis

Statistical analysis on the various *E. diaphana* groups was performed in MetaboAnalyst 5.0 and the normalized data was log transformed with no scaling applied. Data normality and homogeneity were visually confirmed. All 631 peaks were used to generate PCAs and a one-way ANOVA was used to test for differences among all groups (B1-anemones, C1^acro^-anemones, aposymbiotic anemones). The top 60 most significantly different peaks were visualized with heatmaps generated using Euclidean distance and Ward clustering algorithm. The data were subset into pairs of interest (aposymbiotic vs symbiotic anemones; B1-anemones vs C1^acro^-anemones) and t-tested, with the p values corrected by the Benjamini–Hochberg method (Benjamini & Hochberg, 1995). A peak was considered significant when the Padj < 0.05 and fold change (FC) > 1.3, where FC was calculated based on the non-log transformed data. Metabolites found to be significant were spatially visualized and confirmed as either a host component based on its correlation with the host tissue distribution and its presence in the aposymbiotic sample, or as an algal component based on correlation with symbiont distribution and detection in the TLC analysis of Symbiodiniaceae cultures. For *G. fascicularis*, the spatial distribution of specific metabolites of interest were visualized in SCiLS, but no statistical comparison was made.

## Acknowledgements

We thank S. J. T Min Ching and R. Sakamoto for anemone supply; the National Sea Simulator team, especially C. Thompson and L. Koukoumaftsis for coral collection; Metabolomics Australia staffs, especially V. Lui and V. Narayana for fruitful discussion and MSI technical support. This research was supported by the Australian Research Council Laureate Fellowship to MJHvO (FL180100036), the Paul G. Allen Family Foundation, and the Reef Restoration and Adaptation Program, which is funded by the partnership between the Australian Governments Reef Trust and the Great Barrier Reef Foundation.

## Author contributions

Conceptualization: MJHvO, DR, WYC; Methodology: DR, WYC; Investigation: WYC, DR; Visualization: WYC, DR; Writing—original draft: WYC, MJHvO; Writing—review & editing: MJHvO, WYC, DR.

## Conflict of interests

The authors declare that they have no competing interests.

## Data availability

All data are available in the main text or the supplementary materials. Surface area normalized metabolite relative intensities are supplied as Supplementary dataset S1. Raw spectra files for anemones (Supplementary dataset S2-3) and corals (Supplementary dataset S4-5) are available in: https://cloudstor.aarnet.edu.au/plus/s/idLfJ1Zuwqfxnhd. Additional information related to this paper can be requested from the authors.

## Notes

### Competing Interest Statement

The authors have declared no competing interest.

