## Supplementary materials for "Spatial metabolomics for symbiotic marine invertebrates"

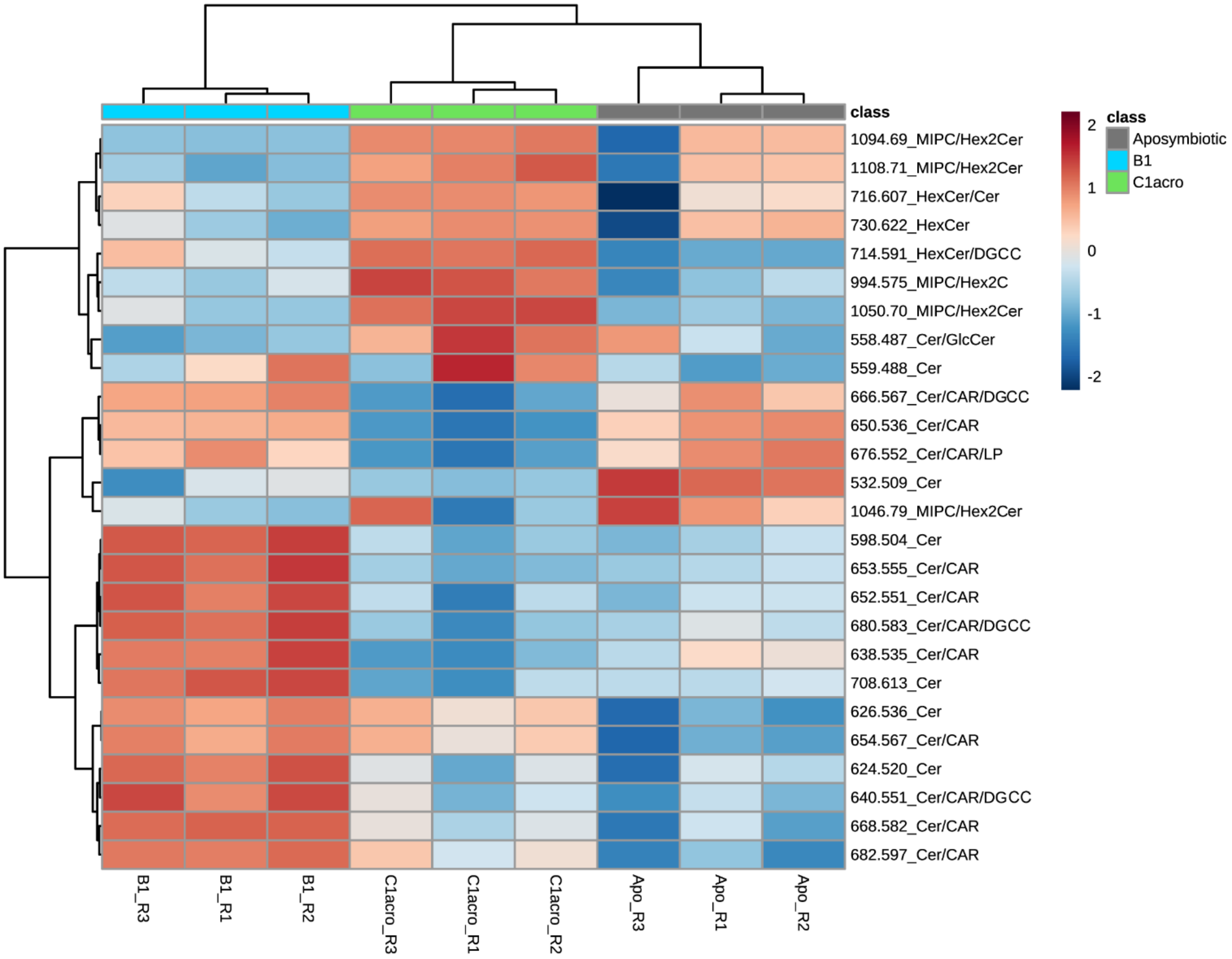


**Key metabolites**

| | |
|---|---|
| Cer | Ceremide |
| HexCer | Hexosylceramide |
| MIPC | Mannosylinositol phosphorylceramide |
| CAR | Acyl carnitine |
| DG | Diglyceride |

**Fig. S1. Heatmap of ceramides in the anemone groups**. Relative intensity of ceramides (26 of the major chemical species detected) in the anemones. Abbreviations of the anemone groups refer to the algal symbionts they were in symbiosis with: B1: *B. minutum*, C1$^{acro}$: *Cladocopium* C1$^{acro}$. "R" in the sample name refers the replicate number.

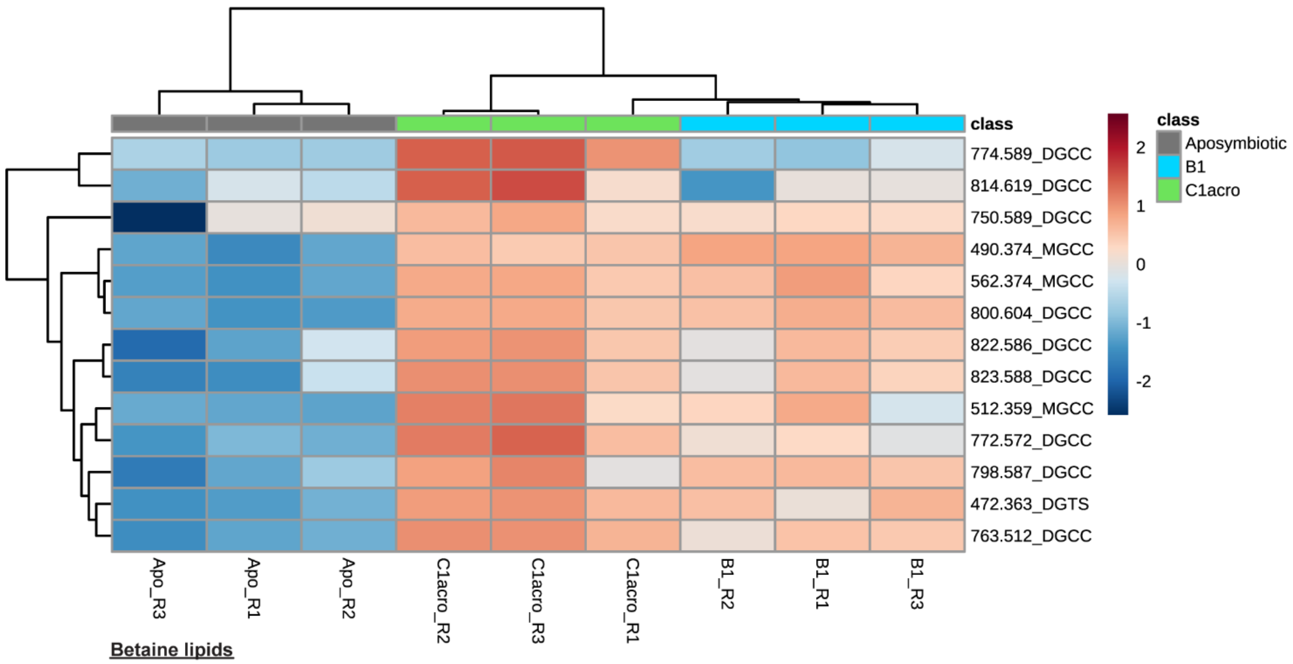


**Betaine lipids**

| | |
|---|---|
| DGCC | Diacylglyceryl-carboxyhydroxy-methylcholine |
| DGTS | Diacylglyceryl-trimethylhomoserine |
| MGCC | Monoacylglyceryl-carboxyhydroxy-methylcholine |

**Fig. S2. Heatmap of betaine lipids in the anemone groups.** Relative intensity of betaine lipids (13 of the major chemical species detected) in aposymbiotic anemones, anemones in symbiosis with the homologous B1 (*B. minutum*) and anemones in symbiosis with the heterologous C1$^{acro}$. "R" in the sample name refers the replicate number. See Table S2 for saturation state of the betaine lipids.

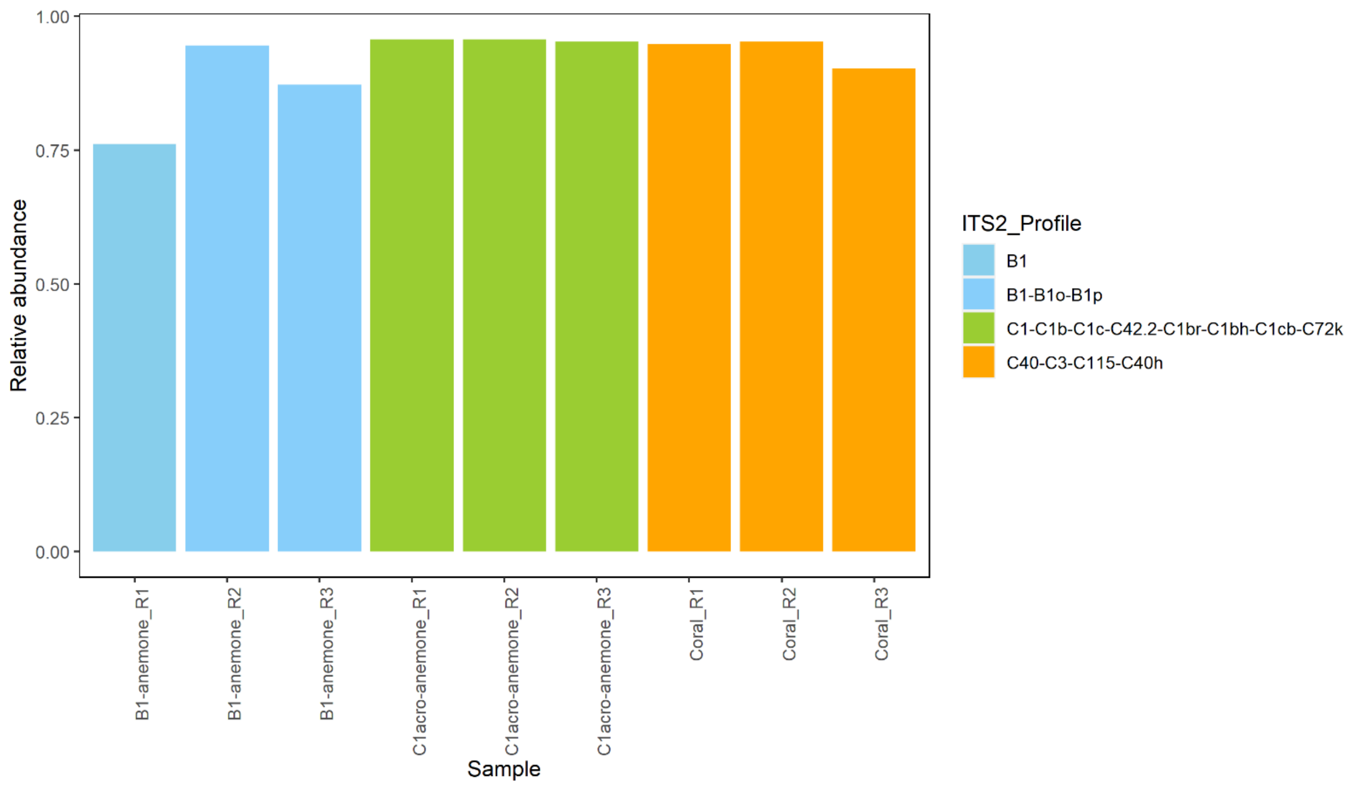


**Fig. S3. Symbiodiniaceae symbiont community (ITS2 profile) of the anemones and corals used in this study.** "R" in the sample name refers the replicate number. Note that the relative abundances do not add up to 1 (i.e., 100%) as some sequences have no ITS2 profile. Anemone data were extracted from Tsang Min Ching et al. (2022)

Reference: Tsang Min Ching SJ, Chan WY, Perez-Gonzalez A, Hillyer KE, Buerger P & van Oppen MJH (2022) Colonization and metabolite profiles of homologous, heterologous and experimentally evolved algal symbionts in the sea anemone *Exaiptasia diaphana*. *ISME COMMUN* 2: 1–10

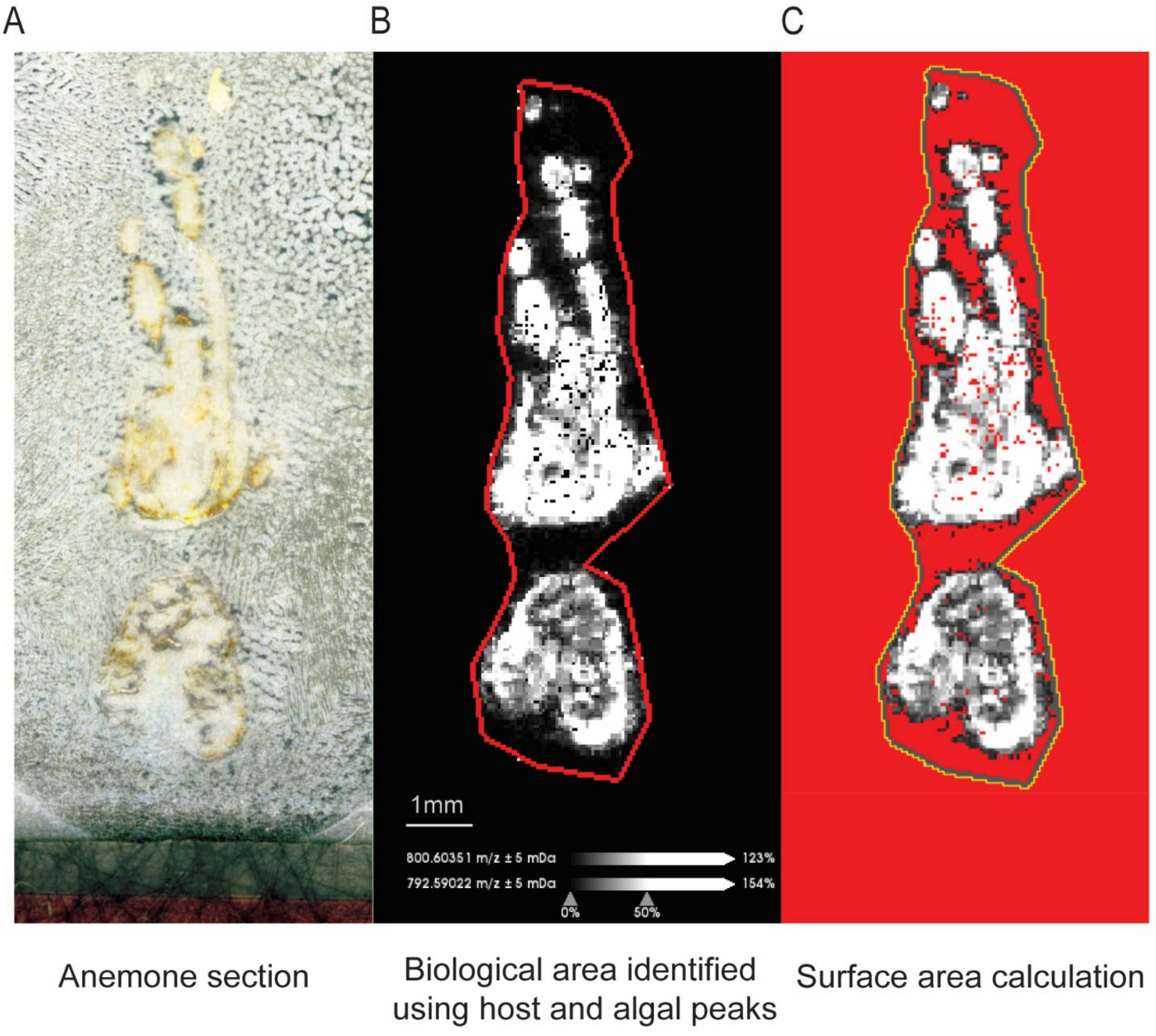


**Fig. S4. Procedures to calculate surface area for ion intensity normalization.** (**A**) image of an anemone section, (**B**) biological area of the tissue section on SCiLS, identified by overlaying the ion density maps of the most prominent peaks associated with the anemone host (m/z 792.590) and the algal symbionts (m/z 800.604), (**C**) surface area identified on ImageJ using the SCiLS exported image under the threshold of 30 for area calculation.

**Table S1. Full names of metabolite abbreviations of the major groups.**

| Abbreviation | Full name | Category |
|---|---|---|
| CAR | Acyl carnitine | Energy homeostasis |
| Cer | Ceramide | Signaling |
| Chl F | Chlorophyll fragment | Energy/photosynthesis |
| DG | Diglyceride | Energy |
| DGCC | Diacylglycerylcarboxyhydroxymethylcholine | Structural |
| DGDG | Digalactosyldiacylglycerol | Structural |
| DGTA | Diacylglycerylhydroxymethyltrimethylalanine | Structural |
| DGTS | Diacylglyceryltrimethylhomoserine | Structural |
| FA | Fatty acids/esters | Energy/backbone |
| FA CONJ | Fatty acid conjugation | Energy/signaling |
| GlcCer | Glucosylceramide | Signaling/immune/cellular recognition |
| HexCer | Hexosylceramide | Signaling/immune/cellular recognition |
| HG | Headgroup | Structural |
| LPA | Lysophosphatidic acid | Signaling |
| LPC | Lysophosphatidylcholine | Structural |
| LPI | Lysophosphatidylinositol | Structural |
| MG | Mono(acyl/alkyl)glycerol | Energy |
| MGCC | Monoacylglycerylcarboxyhydroxymethylcholine | Structural |
| MIPC | Mannosylinositol phosphorylceramide | Signaling/immune/cellular recognition |
| PA | Phosphatidic acid | Signaling |
| PC | Glycerophosphocholine | Structural |
| PE | Phosphatidylethanolamine | Structural |
| PG | Phosphoglycan | Structural/immune |
| Pheo a | Pheophorbide a | Energy/photosynthesis |
| PI | Phosphatidylinositol | Structural/signaling |
| PIP | Phosphatidylinositol-*n*-phosphate | Structural/signaling |
| PM | Polar metabolite | All |
| PS | Phosphatidylserine | Structural/signaling |
| SQDG | Sulfoquinovosyl diacylglycerol | Structural/energy/photosynthesis |
| ST | Sterol | Structural/signaling |
| TG | Triglyceride | Energy |

**Table S2. Ceramides with significantly different relative intensity between C1$^{acro}$-anemones and B1-anemones.**

| m/z | t.stat | $P_{adj}$ | FC* | log2(FC) |
|---|---|---|---|---|
| 558.487_Cer/GlcCer | -6.8 | 0.005 | 0.09 | -3.51 |
| 598.504_Cer | 9.8 | 0.002 | 14.27 | 3.84 |
| 624.520_Cer | 5.0 | 0.01 | 6.18 | 2.63 |
| 638.535_Cer/CAR | 11.4 | 0.002 | 13.30 | 3.73 |
| 640.551_Cer/CAR/DGTS | 5.1 | 0.01 | 5.35 | 2.42 |
| 650.536_Cer/CAR | 16.1 | 0.001 | 10.91 | 3.45 |
| 652.551_Cer/CAR | 5.3 | 0.01 | 6.42 | 2.68 |
| 653.555_Cer/CAR | 12.8 | 0.002 | 13.48 | 3.75 |
| 666.567_Cer/CAR/DG | 10.7 | 0.002 | 11.72 | 3.55 |
| 668.582_Cer/CAR | 8.5 | 0.003 | 4.34 | 2.12 |
| 676.552_Cer/CAR/LPC | 8.0 | 0.003 | 11.17 | 3.48 |
| 680.583_Cer/CAR/DGTS | 9.1 | 0.002 | 10.69 | 3.42 |
| 682.597_Cer/CAR | 5.1 | 0.01 | 7.25 | 2.86 |
| 708.613_Cer | 7.7 | 0.003 | 20.74 | 4.37 |
| 714.591_HexCer/DGCC | -4.3 | 0.017 | 0.17 | -2.58 |
| 716.607_HexCer/Cer | -3.8 | 0.023 | 0.27 | -1.90 |
| 730.622_HexCer | -5.6 | 0.009 | 0.19 | -2.43 |
| 994.575_MIPC/Hex2Cer | -9.2 | 0.002 | 0.18 | -2.47 |
| 1050.70_MIPC/Hex2Cer | -8.0 | 0.003 | 0.05 | -4.39 |
| 1094.69_MIPC/Hex2Cer | -45.4 | <0.001 | 0.03 | -5.21 |
| 1108.71_MIPC/Hex2Cer | -9.4 | 0.002 | 0.05 | -4.39 |

*A fold change (FC) > 1 suggests that this ceramide had higher relative intensity in B1-anemones, whereas a FC < 1 indicates that this metabolite had higher relative intensity in C1$^{acro}$-anemones. For instance, a FC value of 0.1 means that this ceramide had 10 times high relative intensity in C1$^{acro}$-anemones than B1-anemones.

**Table S3. ANOVA and Tukey's HSD results of betaine lipids.** Anemone groups for comparisons were B1-anemones (B1), C1$^{acro}$-anemones (C1$^{acro}$) and aposymbiotic anemones.

| m/z | f.value | FDR | Significant pairs |
|---|---|---|---|
| 472.363_DGTS | 60.4 | <0.001 | B1-Aposymbiotic; C1$^{acro}$-Aposymbiotic |
| 490.374_MGCC | 229.4 | <0.001 | B1-Aposymbiotic; C1$^{acro}$-Aposymbiotic |
| 512.359_MGCC | 18.9 | 0.004 | B1-Aposymbiotic; C1$^{acro}$-Aposymbiotic |
| 562.374_MGCC | 87.0 | <0.001 | B1-Aposymbiotic; C1$^{acro}$-Aposymbiotic |
| 763.512_DGCC | 80.4 | <0.001 | B1-Aposymbiotic; C1$^{acro}$-Aposymbiotic; C1acro-B1 |
| 772.572_DGCC | 43.5 | <0.001 | B1-Aposymbiotic; C1$^{acro}$-Aposymbiotic; C1acro-B1 |
| 774.589_DGCC | 61.7 | <0.001 | C1$^{acro}$-Aposymbiotic; C1acro-B1 |
| 798.587_DGCC | 16.5 | 0.005 | B1-Aposymbiotic; C1$^{acro}$-Aposymbiotic |
| 800.604_DGCC | 228.5 | <0.001 | B1-Aposymbiotic; C1$^{acro}$-Aposymbiotic |
| 822.586_DGCC | 10.5 | 0.013 | B1-Aposymbiotic; C1$^{acro}$-Aposymbiotic |
| 823.588_DGCC | 14.1 | 0.007 | B1-Aposymbiotic; C1$^{acro}$-Aposymbiotic |

**Table S4. T-test results and fold changes of metabolites with significantly different relative intensity between aposymbiotic and symbiotic anemones.**

| m/z | t.stat | $P_{adj}$ | FC* | log2(FC) |
|---|---|---|---|---|
| 383.288_FA/ST | 3.0 | 0.050 | 11.93 | 3.58 |
| 397.303_FA | 3.5 | 0.032 | 27.45 | 4.78 |
| 465.353_FA CONJ | -4.0 | 0.020 | 0.02 | -5.39 |
| 472.363_DGTS | -9.1 | 0.001 | 0.02 | -5.97 |
| 490.374_MGCC | -15.8 | <0.001 | 0.002 | -9.32 |
| 512.359_MGCC | -5.3 | 0.006 | 0.04 | -4.72 |
| 532.509_Cer | 6.8 | 0.002 | 38.07 | 5.25 |
| 562.374_MGCC | -14.0 | <0.001 | 0.002 | -8.72 |
| 592.268_Pheo a | -3.3 | 0.039 | 0.07 | -3.77 |
| 593.276_Pheo a | -3.4 | 0.035 | 0.05 | -4.24 |
| 614.427_Chl F | -3.7 | 0.027 | 0.06 | -3.94 |
| 625.507_DG | 5.1 | 0.007 | 22.41 | 4.49 |
| 626.536_Cer | -7.6 | 0.002 | 0.02 | -5.44 |
| 641.352_PM | 3.1 | 0.045 | 5.60 | 2.49 |
| 654.567_Cer/CAR | -6.8 | 0.002 | 0.04 | -4.54 |
| 655.570_DG | -5.2 | 0.007 | 0.03 | -5.16 |
| 661.515_LPA | 3.5 | 0.030 | 22.78 | 4.51 |
| 682.597_Cer/CAR | -4.7 | 0.010 | 0.02 | -5.88 |
| 693.373_PG/LPG | -3.8 | 0.024 | 0.12 | -3.07 |
| 714.591_HexCer/DGCC | -4.1 | 0.017 | 0.03 | -5.01 |
| 717.577_DG | 3.8 | 0.024 | 27.77 | 4.80 |
| 732.503_PC | -3.8 | 0.024 | 0.12 | -3.03 |
| 732.554_PC/PE | 5.5 | 0.005 | 9.19 | 3.20 |
| 733.516_PA | 3.4 | 0.033 | 7.68 | 2.94 |
| 734.556_DGCC | 3.2 | 0.044 | 11.13 | 3.48 |
| 740.559_PC | 8.9 | 0.001 | 15.05 | 3.91 |
| 741.561_DG/TG | 3.1 | 0.045 | 6.43 | 2.69 |
| 744.590_PC | 9.8 | 0.001 | 7.83 | 2.97 |
| 745.479_PG | 3.1 | 0.048 | 6.51 | 2.70 |
| 746.461_DGCC | -3.4 | 0.035 | 0.09 | -3.49 |
| 746.605_PC/PE | 3.7 | 0.027 | 3.55 | 1.83 |
| 747.609_DG/TG | 4.4 | 0.013 | 4.29 | 2.10 |
| 748.572_DGCC | -3.4 | 0.033 | 0.04 | -4.70 |
| 749.576_DG/TG | -3.5 | 0.030 | 0.05 | -4.36 |
| 756.590_PC | 5.1 | 0.007 | 25.22 | 4.66 |
| 759.573_PC/PE | 8.2 | 0.001 | 7.49 | 2.90 |
| 760.534_PC/PE | -4.4 | 0.013 | 0.15 | -2.78 |

| | | | | |
|---|---|---|---|---|
| 760.585_PC/PE | 5.3 | 0.006 | 5.63 | 2.49 |
| 761.588_PC/PE | 5.0 | 0.008 | 6.28 | 2.65 |
| 763.512_DGCC | -8.2 | 0.001 | 0.005 | -7.70 |
| 764.558_PC | 4.3 | 0.014 | 3.92 | 1.97 |
| 765.560_PC/PE | 4.2 | 0.015 | 6.13 | 2.62 |
| 768.591_PC | 3.1 | 0.045 | 2.33 | 1.22 |
| 769.593_PC | 3.7 | 0.027 | 2.58 | 1.37 |
| 770.605_PC | 11.2 | <0.001 | 27.02 | 4.76 |
| 772.572_DGCC | -4.7 | 0.010 | 0.03 | -5.21 |
| 773.171_PM | -3.1 | 0.047 | 0.01 | -7.17 |
| 779.486_PG | -6.4 | 0.003 | 0.02 | -5.38 |
| 782.568_PC | 10.8 | <0.001 | 11.62 | 3.54 |
| 784.534_PC | -5.0 | 0.008 | 0.11 | -3.21 |
| 784.584_PC | 9.8 | 0.001 | 22.26 | 4.48 |
| 785.453_SQDG | -7.1 | 0.002 | 0.03 | -4.98 |
| 787.604_PC | 8.8 | 0.001 | 43.15 | 5.43 |
| 789.620_PA/TG | 5.7 | 0.005 | 26.94 | 4.75 |
| 796.622_PC | 3.3 | 0.036 | 3.23 | 1.69 |
| 798.587_DGCC | -6.2 | 0.003 | 0.05 | -4.20 |
| 800.604_DGCC | -22.8 | <0.001 | 0.05 | -9.35 |
| 802.536_PC | 3.5 | 0.030 | 7.29 | 2.87 |
| 806.567_PC | 5.9 | 0.004 | 5.27 | 2.40 |
| 807.571_PC | 6.7 | 0.002 | 6.35 | 2.67 |
| 808.584_PC | 7.1 | 0.002 | 8.37 | 3.07 |
| 809.587_PC | 7.7 | 0.002 | 11.57 | 3.53 |
| 810.600_PC | 5.6 | 0.005 | 7.62 | 2.93 |
| 811.604_PC | 5.4 | 0.006 | 10.95 | 3.45 |
| 822.586_DGCC | -4.4 | 0.013 | 0.07 | -3.93 |
| 822.636_PC | 5.4 | 0.006 | 39.27 | 5.30 |
| 823.588_DGCC | -4.8 | 0.009 | 0.04 | -4.73 |
| 834.601_PC | 3.5 | 0.030 | 6.23 | 2.64 |
| 835.603_PC | 3.4 | 0.035 | 8.40 | 3.07 |
| 836.615_PC | 4.9 | 0.008 | 13.13 | 3.71 |
| 838.633_PC | 3.9 | 0.020 | 6.37 | 2.67 |
| 839.563_Chl | -6.6 | 0.003 | 0.04 | -4.74 |
| 839.635_PC | 4.1 | 0.018 | 7.45 | 2.90 |
| 870.564_PC | -6.5 | 0.003 | 0.02 | -5.84 |
| 988.686_PC | -3.2 | 0.040 | 0.05 | -4.42 |
| 1019.61_PIP/DGDG | 3.4 | 0.033 | 5.48 | 2.45 |
| 400.302 | -12.1 | <0.001 | 0.01 | -6.43 |

| | | | | |
|---|---|---|---|---|
| 400.804 | -6.9 | 0.002 | 0.03 | -5.25 |
| 464.35 | -4.8 | 0.009 | 0.09 | -3.44 |
| 491.377 | -26.1 | <0.001 | 0.01 | -6.87 |
| 496.376 | 6.5 | 0.003 | 7.22 | 2.85 |
| 497.38 | 5.3 | 0.006 | 9.16 | 3.20 |
| 500.483 | 7.5 | 0.002 | 23.38 | 4.55 |
| 508.377 | 5.1 | 0.007 | 36.62 | 5.19 |
| 518.493 | 5.8 | 0.005 | 8.91 | 3.16 |
| 519.497 | 5.2 | 0.007 | 16.64 | 4.06 |
| 522.355 | 4.7 | 0.010 | 15.68 | 3.97 |
| 524.145 | 4.0 | 0.020 | 24.29 | 4.60 |
| 524.371 | 6.9 | 0.002 | 9.71 | 3.28 |
| 524.407 | 6.1 | 0.004 | 6.53 | 2.71 |
| 525.374 | 4.5 | 0.011 | 14.46 | 3.85 |
| 547.357 | 5.1 | 0.007 | 8.59 | 3.10 |
| 562.158 | 4.5 | 0.012 | 12.50 | 3.64 |
| 563.378 | -11.3 | <0.001 | 0.003 | -8.35 |
| 564.381 | -3.7 | 0.025 | 0.06 | -3.99 |
| 568.135 | 4.1 | 0.018 | 28.37 | 4.83 |
| 578.369 | -3.8 | 0.023 | 0.05 | -4.40 |
| 581.399 | -8.4 | 0.001 | 0.01 | -6.74 |
| 582.402 | -5.3 | 0.006 | 0.02 | -5.32 |
| 584.356 | -3.7 | 0.027 | 0.04 | -4.75 |
| 590.322 | -4.5 | 0.012 | 0.01 | -6.14 |
| 592.035 | 3.4 | 0.033 | 7.46 | 2.90 |
| 602.427 | -4.5 | 0.012 | 0.04 | -4.56 |
| 606.296 | -4.0 | 0.020 | 0.01 | -6.30 |
| 612.519 | -3.3 | 0.037 | 0.05 | -4.47 |
| 621.421 | -3.1 | 0.048 | 0.22 | -2.19 |
| 622.026 | -7.4 | 0.002 | 0.02 | -5.62 |
| 622.084 | -3.6 | 0.027 | 0.05 | -4.24 |
| 626.427 | -5.1 | 0.007 | 0.05 | -4.22 |
| 636.223 | -3.4 | 0.035 | 0.03 | -5.15 |
| 643.517 | 4.0 | 0.019 | 6.07 | 2.60 |
| 644.453 | -3.6 | 0.028 | 0.03 | -5.11 |
| 662.518 | 3.9 | 0.021 | 20.74 | 4.37 |
| 665.5 | 7.1 | 0.002 | 3.51 | 1.81 |
| 666.504 | 5.1 | 0.007 | 4.05 | 2.02 |
| 667.506 | 5.0 | 0.008 | 9.79 | 3.29 |
| 674.163 | -13.6 | <0.001 | 0.02 | -5.66 |

| | | | | |
|---|---|---|---|---|
| 679.457 | -3.8 | 0.024 | 0.30 | -1.74 |
| 680.519 | 6.8 | 0.002 | 16.08 | 4.01 |
| 742.575 | 12.8 | <0.001 | 7.77 | 2.96 |
| 743.578 | 11.3 | <0.001 | 9.21 | 3.20 |
| 744.493 | -3.4 | 0.034 | 0.22 | -2.21 |
| 745.495 | -4.7 | 0.010 | 0.12 | -3.00 |
| 745.593 | 10.6 | <0.001 | 10.78 | 3.43 |
| 748.477 | -3.5 | 0.033 | 0.14 | -2.86 |
| 748.612 | 3.0 | 0.050 | 5.23 | 2.39 |
| 751.592 | -3.2 | 0.040 | 0.15 | -2.73 |
| 752.603 | -8.8 | 0.001 | 0.05 | -4.31 |
| 754.536 | 10.5 | <0.001 | 28.06 | 4.81 |
| 754.619 | -12.8 | <0.001 | 0.02 | -6.05 |
| 755.498 | -15.5 | <0.001 | 0.02 | -5.91 |
| 755.623 | -6.0 | 0.004 | 0.05 | -4.42 |
| 756.503 | -10.3 | <0.001 | 0.01 | -6.65 |
| 756.554 | 8.6 | 0.001 | 5.95 | 2.57 |
| 757.557 | 7.6 | 0.002 | 10.84 | 3.44 |
| 758.569 | 5.2 | 0.007 | 6.03 | 2.59 |
| 760.622 | 6.7 | 0.003 | 11.19 | 3.48 |
| 762.59 | 3.8 | 0.024 | 8.02 | 3.00 |
| 764.516 | -5.8 | 0.004 | 0.02 | -5.97 |
| 768.577 | 3.1 | 0.047 | 2.53 | 1.34 |
| 769.478 | -5.9 | 0.004 | 0.05 | -4.38 |
| 770.482 | -6.1 | 0.004 | 0.04 | -4.57 |
| 771.473 | -12.4 | <0.001 | 0.01 | -6.88 |
| 771.609 | 9.7 | 0.001 | 30.03 | 4.91 |
| 772.477 | -13.4 | <0.001 | 0.02 | -5.93 |
| 772.525 | 4.5 | 0.012 | 4.42 | 2.14 |
| 772.621 | 10.7 | <0.001 | 23.38 | 4.55 |
| 773.575 | -3.7 | 0.026 | 0.06 | -3.95 |
| 773.624 | 8.8 | 0.001 | 63.33 | 5.98 |
| 774.601 | 4.3 | 0.014 | 12.10 | 3.60 |
| 774.636 | 4.2 | 0.015 | 10.75 | 3.43 |
| 775.64 | 6.8 | 0.002 | 10.81 | 3.43 |
| 778.537 | 3.1 | 0.046 | 2.80 | 1.49 |
| 779.363 | -3.0 | 0.048 | 0.05 | -4.45 |
| 780.552 | 7.2 | 0.002 | 5.25 | 2.39 |
| 780.589 | 3.6 | 0.027 | 5.16 | 2.37 |
| 781.556 | 5.8 | 0.005 | 8.73 | 3.13 |

| | | | | |
|---|---|---|---|---|
| 781.593 | 3.9 | 0.020 | 7.47 | 2.90 |
| 783.53 | -7.5 | 0.002 | 0.05 | -4.37 |
| 783.571 | 11.0 | <0.001 | 11.18 | 3.48 |
| 785.588 | 8.0 | 0.001 | 27.49 | 4.78 |
| 786.601 | 9.7 | 0.001 | 21.02 | 4.39 |
| 788.615 | 6.1 | 0.004 | 17.09 | 4.09 |
| 794.607 | 5.4 | 0.006 | 5.54 | 2.47 |
| 795.608 | 4.9 | 0.009 | 7.01 | 2.81 |
| 797.624 | 3.3 | 0.037 | 3.43 | 1.78 |
| 798.628 | 8.9 | 0.001 | 11.01 | 3.46 |
| 801.606 | -16.3 | <0.001 | 0.001 | -9.78 |
| 802.611 | -14.9 | <0.001 | 0.002 | -8.83 |
| 803.613 | -10.1 | <0.001 | 0.02 | -5.94 |
| 804.552 | 6.3 | 0.003 | 11.66 | 3.54 |
| 805.554 | 5.1 | 0.007 | 15.02 | 3.91 |
| 808.562 | 3.8 | 0.024 | 12.97 | 3.70 |
| 808.621 | 4.1 | 0.018 | 8.26 | 3.05 |
| 810.451 | -5.6 | 0.005 | 0.02 | -5.82 |
| 810.464 | -3.2 | 0.040 | 0.17 | -2.58 |
| 812.616 | 8.5 | 0.001 | 58.00 | 5.86 |
| 814.571 | -10.3 | <0.001 | 0.13 | -2.96 |
| 815.575 | -10.5 | <0.001 | 0.07 | -3.88 |
| 815.622 | -4.6 | 0.011 | 0.01 | -6.29 |
| 816.588 | 3.6 | 0.028 | 2.73 | 1.45 |
| 816.597 | -8.2 | 0.001 | 0.01 | -6.10 |
| 817.592 | 3.1 | 0.046 | 2.93 | 1.55 |
| 820.526 | 3.2 | 0.044 | 13.47 | 3.75 |
| 822.541 | 3.0 | 0.050 | 9.90 | 3.31 |
| 828.551 | -4.5 | 0.012 | 0.28 | -1.85 |
| 828.635 | -10.5 | <0.001 | 0.02 | -5.86 |
| 829.556 | -4.8 | 0.009 | 0.24 | -2.04 |
| 830.546 | -4.2 | 0.016 | 0.08 | -3.59 |
| 831.549 | -7.5 | 0.002 | 0.03 | -5.24 |
| 832.583 | 5.6 | 0.005 | 4.89 | 2.29 |
| 832.593 | -6.3 | 0.003 | 0.03 | -5.22 |
| 833.587 | 5.7 | 0.005 | 6.12 | 2.61 |
| 838.559 | -5.0 | 0.008 | 0.12 | -3.04 |
| 842.604 | -3.1 | 0.046 | 0.36 | -1.47 |
| 843.607 | -4.6 | 0.011 | 0.19 | -2.43 |
| 845.529 | -3.6 | 0.028 | 0.13 | -2.97 |

| | | | | |
|---|---|---|---|---|
| 854.235 | -3.1 | 0.045 | 0.01 | -7.31 |
| 854.244 | -3.3 | 0.037 | 0.01 | -7.60 |
| 854.543 | 5.1 | 0.007 | 8.09 | 3.02 |
| 859.579 | -3.1 | 0.047 | 0.15 | -2.71 |
| 864.585 | -5.7 | 0.005 | 0.03 | -5.28 |
| 866.681 | -5.3 | 0.007 | 0.05 | -4.37 |
| 867.683 | -5.2 | 0.007 | 0.03 | -5.25 |
| 871.571 | -6.5 | 0.003 | 0.01 | -7.63 |
| 872.577 | -5.0 | 0.008 | 0.02 | -5.97 |
| 872.603 | -7.0 | 0.002 | 0.02 | -5.98 |
| 873.607 | -5.2 | 0.007 | 0.03 | -5.03 |
| 878.816 | -3.1 | 0.045 | 0.04 | -4.59 |
| 880.833 | -3.4 | 0.035 | 0.02 | -6.02 |
| 885.646 | -6.5 | 0.003 | 0.04 | -4.81 |
| 893.503 | -3.3 | 0.036 | 0.24 | -2.05 |
| 893.71 | -3.7 | 0.025 | 0.18 | -2.48 |
| 899.406 | -3.3 | 0.037 | 0.18 | -2.51 |
| 906.847 | -3.5 | 0.030 | 0.02 | -5.68 |
| 907.85 | -3.6 | 0.027 | 0.03 | -5.09 |
| 910.714 | -12.6 | <0.001 | 0.005 | -7.69 |
| 911.716 | -8.7 | 0.001 | 0.01 | -6.53 |
| 912.72 | -6.1 | 0.004 | 0.03 | -5.30 |
| 920.385 | -4.1 | 0.018 | 0.09 | -3.49 |
| 930.849 | -3.1 | 0.045 | 0.03 | -5.23 |
| 932.866 | -4.4 | 0.013 | 0.03 | -5.26 |
| 954.603 | 3.4 | 0.035 | 5.91 | 2.56 |
| 981.63 | 6.6 | 0.003 | 9.98 | 3.32 |
| 982.634 | 7.4 | 0.002 | 13.28 | 3.73 |
| 983.646 | 10.1 | <0.001 | 20.38 | 4.35 |
| 985.662 | 4.6 | 0.011 | 5.61 | 2.49 |
| 989.644 | -4.3 | 0.014 | 0.05 | -4.27 |
| 993.592 | 4.3 | 0.014 | 10.99 | 3.46 |
| 994.598 | 4.6 | 0.011 | 12.74 | 3.67 |
| 995.61 | 5.6 | 0.005 | 10.29 | 3.36 |
| 996.613 | 5.2 | 0.007 | 12.35 | 3.63 |
| 997.627 | 6.3 | 0.003 | 16.26 | 4.02 |
| 999.642 | 4.9 | 0.008 | 10.47 | 3.39 |
| 1003.61 | -9.6 | 0.001 | 0.22 | -2.19 |
| 1004.62 | -7.1 | 0.002 | 0.18 | -2.50 |
| 1005.62 | -5.3 | 0.006 | 0.11 | -3.17 |

| | | | | |
|---|---|---|---|---|
| 1005.63 | 3.8 | 0.024 | 3.88 | 1.96 |
| 1019.59 | -5.6 | 0.005 | 0.07 | -3.76 |
| 1020.59 | -6.2 | 0.003 | 0.05 | -4.34 |
| 1021.62 | 3.3 | 0.037 | 5.55 | 2.47 |
| 1022.63 | 3.7 | 0.025 | 6.48 | 2.70 |
| 1043.58 | 5.5 | 0.005 | 14.31 | 3.84 |
| 1044.59 | 5.8 | 0.005 | 15.95 | 4.00 |
| 1054.72 | -4.7 | 0.010 | 0.12 | -3.02 |
| 1055.72 | -3.5 | 0.032 | 0.17 | -2.57 |
| 1146.68 | 6.9 | 0.002 | 7.29 | 2.87 |
| 1160.65 | 3.4 | 0.035 | 5.69 | 2.51 |
| 1215.63 | -3.0 | 0.050 | 0.09 | -3.50 |

*A fold change (FC) > 1 suggests that this metabolite had higher relative intensity in aposymbiotic anemones, whereas a FC < 1 indicates that this metabolite had higher relative intensity in symbiotic anemones. For instance, a FC value of 0.1 means that this metabolite had 10 times high relative intensity in symbiotic than aposymbiotic anemones.

**Table S5. T-test results and fold changes of metabolites with significantly different relative intensity between B1-anemoens and C1$^{acro}$-anemones.**

| m/z | t.stat | $P_{adj}$ | FC* | log2(FC) |
|---|---|---|---|---|
| 184.073_PC HG | 9.6 | 0.012 | 2839.70 | 11.47 |
| 241.180_FA | 6.9 | 0.017 | 113.10 | 6.82 |
| 256.203_FA | 5.6 | 0.027 | 116.01 | 6.86 |
| 264.204_FA/MG | 8.5 | 0.014 | 126.40 | 6.98 |
| 336.225_CAR | -9.5 | 0.012 | 0.002 | -9.02 |
| 337.229_CAR | -18.7 | 0.012 | 0.001 | -9.49 |
| 358.304_CAR/FA | -8.3 | 0.014 | 0.04 | -4.52 |
| 397.303_FA | 5.9 | 0.025 | 47.79 | 5.58 |
| 402.198_FA CONJ | -11.6 | 0.012 | 0.01 | -6.60 |
| 452.181_FA CONJ | -9.7 | 0.012 | 0.06 | -4.17 |
| 465.353_FA CONJ | -4.3 | 0.047 | 0.12 | -3.02 |
| 482.284_Chl F | -5.7 | 0.027 | 0.05 | -4.25 |
| 526.274_Lyso-lipid | -7.1 | 0.017 | 0.04 | -4.78 |
| 545.239_ST | -5.3 | 0.029 | 0.05 | -4.42 |
| 558.487_Cer/GlcCer | -6.8 | 0.018 | 0.09 | -3.51 |
| 573.488_WE/DG | -6.5 | 0.020 | 0.08 | -3.70 |
| 575.226_LPC | -6.8 | 0.018 | 0.03 | -5.13 |
| 597.235_LPI | -9.8 | 0.012 | 0.02 | -5.62 |
| 598.504_Cer | 9.8 | 0.012 | 14.27 | 3.84 |
| 609.524_DG | -5.3 | 0.030 | 0.18 | -2.48 |
| 624.520_Cer | 5.0 | 0.033 | 6.18 | 2.63 |
| 635.233_PM | -5.0 | 0.034 | 0.22 | -2.17 |
| 636.236_PM | -11.9 | 0.012 | 0.12 | -3.11 |
| 638.535_Cer/CAR | 11.4 | 0.012 | 13.30 | 3.73 |
| 640.551_Cer/CAR/DGTS | 5.1 | 0.032 | 5.35 | 2.42 |
| 641.225_PM | -7.5 | 0.017 | 0.04 | -4.75 |
| 641.555_DG | 7.9 | 0.015 | 10.44 | 3.38 |
| 650.536_Cer/CAR | 16.1 | 0.012 | 10.91 | 3.45 |
| 652.551_Cer/CAR | 5.3 | 0.030 | 6.42 | 2.68 |
| 653.555_Cer/CAR | 12.8 | 0.012 | 13.48 | 3.75 |
| 655.570_DG | 4.6 | 0.040 | 4.84 | 2.27 |
| 656.573_DG | 8.6 | 0.014 | 8.88 | 3.15 |
| 657.216_PM | -10.6 | 0.012 | 0.26 | -1.94 |
| 666.567_Cer/CAR/DG | 10.7 | 0.012 | 11.72 | 3.55 |
| 668.582_Cer/CAR | 8.5 | 0.014 | 4.34 | 2.12 |
| 676.552_Cer/CAR/LPC | 8.0 | 0.015 | 11.17 | 3.48 |
| 680.183_PM | -4.8 | 0.036 | 0.06 | -4.10 |

| | | | | |
|---|---|---|---|---|
| 680.583_Cer/CAR/DGTS | 9.1 | 0.012 | 10.69 | 3.42 |
| 681.585_DG | 12.5 | 0.012 | 23.93 | 4.58 |
| 682.597_Cer/CAR | 5.1 | 0.032 | 7.25 | 2.86 |
| 693.373_PG/LPG | -5.7 | 0.026 | 0.26 | -1.95 |
| 705.484_PA | -5.9 | 0.025 | 0.15 | -2.76 |
| 706.487_PA | -5.6 | 0.027 | 0.11 | -3.19 |
| 707.196_PM | -6.2 | 0.022 | 0.09 | -3.45 |
| 707.500_PA | -7.2 | 0.017 | 0.10 | -3.37 |
| 708.613_Cer | 7.7 | 0.016 | 20.74 | 4.37 |
| 714.591_HexCer/DGCC | -4.3 | 0.048 | 0.17 | -2.58 |
| 718.613_LPC | -4.3 | 0.048 | 0.14 | -2.83 |
| 719.463_PA | -5.7 | 0.027 | 0.15 | -2.72 |
| 721.457_PG | -5.5 | 0.028 | 0.17 | -2.54 |
| 721.480_PA | -4.7 | 0.037 | 0.13 | -2.97 |
| 728.607_DGCC | -8.0 | 0.015 | 0.05 | -4.19 |
| 729.484_PA | -8.5 | 0.014 | 0.13 | -2.97 |
| 730.622_HexCer | -5.6 | 0.027 | 0.19 | -2.43 |
| 731.499_PA/SM | -5.3 | 0.030 | 0.32 | -1.63 |
| 731.625_DG/TG | -5.4 | 0.028 | 0.14 | -2.81 |
| 732.345_PS | 5.7 | 0.026 | 6.84 | 2.77 |
| 732.503_PC | -9.4 | 0.012 | 0.24 | -2.09 |
| 733.516_PA | -5.9 | 0.025 | 0.12 | -3.01 |
| 742.287_PC | -8.3 | 0.014 | 0.03 | -4.91 |
| 745.479_PG | -4.8 | 0.036 | 0.13 | -2.97 |
| 747.473_SQDG | -4.9 | 0.035 | 0.35 | -1.51 |
| 756.590_PC | -5.2 | 0.031 | 0.14 | -2.81 |
| 758.519_PC/PE | -5.0 | 0.033 | 0.21 | -2.26 |
| 760.534_PC/PE | -4.9 | 0.035 | 0.32 | -1.64 |
| 762.540_PC/PE | -6.1 | 0.024 | 0.12 | -3.10 |
| 764.558_PC | -6.3 | 0.021 | 0.36 | -1.46 |
| 765.560_PC/PE | -7.2 | 0.017 | 0.23 | -2.10 |
| 770.570_PE/PC | -4.7 | 0.037 | 0.12 | -3.09 |
| 773.171_PM | -5.1 | 0.032 | 0.04 | -4.51 |
| 774.589_DGCC | -7.7 | 0.016 | 0.01 | -6.17 |
| 775.592_TG | -5.4 | 0.028 | 0.03 | -4.92 |
| 779.150_PM | -7.2 | 0.017 | 0.04 | -4.83 |
| 781.534_PG/TG | -4.9 | 0.035 | 0.26 | -1.95 |
| 782.546_DGCC | -8.2 | 0.015 | 0.28 | -1.86 |
| 789.561_PC | -4.5 | 0.043 | 0.29 | -1.80 |
| 802.536_PC | -4.6 | 0.038 | 0.13 | -2.91 |

| | | | | |
|---|---|---|---|---|
| 823.631_PC | -9.3 | 0.012 | 0.04 | -4.52 |
| 824.637_PC | -16.5 | 0.012 | 0.03 | -5.25 |
| 825.648_PC | -6.2 | 0.022 | 0.05 | -4.26 |
| 826.651_PC | -5.9 | 0.025 | 0.03 | -4.99 |
| 831.270_PIP | -6.6 | 0.019 | 0.04 | -4.75 |
| 853.681_DG/TG | -10.7 | 0.012 | 0.04 | -4.68 |
| 871.717_TG | -4.8 | 0.035 | 0.17 | -2.58 |
| 913.664_TG | -5.3 | 0.029 | 0.14 | -2.81 |
| 914.667_TG | -6.7 | 0.018 | 0.07 | -3.93 |
| 939.680_TG | -7.4 | 0.017 | 0.09 | -3.50 |
| 953.835_TG | -5.0 | 0.033 | 0.10 | -3.37 |
| 967.711_TG | -6.9 | 0.017 | 0.10 | -3.38 |
| 969.315_PIP2 | -11.7 | 0.012 | 0.03 | -4.93 |
| 988.686_PC | -6.8 | 0.018 | 0.11 | -3.13 |
| 994.575_MIPC/Hex2Cer | -9.2 | 0.012 | 0.18 | -2.47 |
| 1009.57_PI | -5.8 | 0.026 | 0.24 | -2.07 |
| 1019.61_PIP/DGDG | -5.4 | 0.029 | 0.21 | -2.28 |
| 1035.29_PIP | -8.0 | 0.015 | 0.04 | -4.71 |
| 1041.28_PIP | -10.5 | 0.012 | 0.01 | -6.29 |
| 1050.7_MIPC/Hex2Cer | -8.0 | 0.015 | 0.05 | -4.39 |
| 1065.57_PIP/PIP2 | -7.2 | 0.017 | 0.11 | -3.17 |
| 1081.54_PIP/PIP2 | -5.9 | 0.025 | 0.16 | -2.65 |
| 1082.54_PIP/PIP2 | -6.3 | 0.021 | 0.23 | -2.13 |
| 1094.69_MIPC/Hex2Cer | -45.4 | 0.001 | 0.03 | -5.21 |
| 1108.71_MIPC/Hex2Cer | -9.4 | 0.012 | 0.05 | -4.39 |
| 1183.62_PIP2 | -6.1 | 0.024 | 0.09 | -3.42 |
| 356.195 | -13.8 | 0.012 | 0.003 | -8.57 |
| 395.096 | -9.4 | 0.012 | 0.01 | -6.64 |
| 396.797 | 6.2 | 0.023 | 37.61 | 5.23 |
| 408.191 | -7.1 | 0.017 | 0.03 | -4.90 |
| 453.184 | -5.5 | 0.027 | 0.03 | -5.25 |
| 548.256 | -5.1 | 0.032 | 0.09 | -3.48 |
| 569.26 | -11.9 | 0.012 | 0.02 | -5.41 |
| 584.128 | -5.4 | 0.028 | 0.06 | -4.04 |
| 585.525 | -7.0 | 0.017 | 0.05 | -4.31 |
| 591.243 | -8.1 | 0.015 | 0.10 | -3.39 |
| 612.519 | 4.3 | 0.047 | 7.61 | 2.93 |
| 678.567 | 7.5 | 0.016 | 10.44 | 3.38 |
| 683.601 | 6.9 | 0.017 | 17.17 | 4.10 |
| 744.638 | -9.2 | 0.012 | 0.05 | -4.36 |

| | | | | |
|---|---|---|---|---|
| 745.457 | -4.7 | 0.036 | 0.14 | -2.79 |
| 756.503 | -4.9 | 0.035 | 0.37 | -1.44 |
| 779.478 | -10.2 | 0.012 | 0.01 | -6.24 |
| 779.539 | -5.0 | 0.033 | 0.24 | -2.03 |
| 786.277 | -7.5 | 0.016 | 0.02 | -5.51 |
| 787.282 | -9.3 | 0.012 | 0.03 | -4.91 |
| 797.617 | -11.8 | 0.012 | 0.04 | -4.64 |
| 811.129 | -6.0 | 0.025 | 0.08 | -3.71 |
| 817.68 | -5.6 | 0.027 | 0.13 | -2.98 |
| 821.625 | 7.8 | 0.016 | 5.10 | 2.35 |
| 830.268 | -4.5 | 0.041 | 0.05 | -4.35 |
| 850.235 | -4.9 | 0.035 | 0.09 | -3.55 |
| 854.235 | -5.5 | 0.027 | 0.04 | -4.68 |
| 875.235 | -7.0 | 0.017 | 0.04 | -4.77 |
| 876.524 | -7.0 | 0.017 | 0.14 | -2.82 |
| 877.527 | -4.8 | 0.036 | 0.17 | -2.59 |
| 878.816 | -9.3 | 0.012 | 0.08 | -3.56 |
| 880.833 | -9.4 | 0.012 | 0.05 | -4.35 |
| 893.503 | -5.8 | 0.026 | 0.35 | -1.53 |
| 896.238 | -7.1 | 0.017 | 0.04 | -4.70 |
| 902.817 | -10.9 | 0.012 | 0.05 | -4.43 |
| 903.357 | -11.9 | 0.012 | 0.03 | -5.14 |
| 905.651 | -7.5 | 0.016 | 0.07 | -3.81 |
| 906.653 | -10.0 | 0.012 | 0.05 | -4.42 |
| 906.847 | -8.4 | 0.014 | 0.07 | -3.84 |
| 907.85 | -8.4 | 0.014 | 0.09 | -3.44 |
| 912.212 | -6.9 | 0.017 | 0.07 | -3.74 |
| 926.818 | -7.4 | 0.017 | 0.05 | -4.32 |
| 927.632 | -11.6 | 0.012 | 0.04 | -4.58 |
| 928.833 | -8.0 | 0.015 | 0.05 | -4.33 |
| 929.836 | -7.1 | 0.017 | 0.05 | -4.42 |
| 930.849 | -8.1 | 0.015 | 0.06 | -4.06 |
| 932.866 | -6.9 | 0.017 | 0.13 | -2.98 |
| 952.832 | -5.2 | 0.031 | 0.11 | -3.13 |
| 953.599 | -4.8 | 0.035 | 0.18 | -2.46 |
| 954.603 | -4.5 | 0.041 | 0.18 | -2.47 |
| 967.577 | -6.2 | 0.022 | 0.14 | -2.85 |
| 968.58 | -5.2 | 0.030 | 0.13 | -3.00 |
| 969.572 | -4.8 | 0.035 | 0.20 | -2.29 |
| 969.594 | -5.2 | 0.030 | 0.13 | -2.90 |

| | | | | |
|---|---|---|---|---|
| 970.599 | -5.7 | 0.026 | 0.12 | -3.06 |
| 993.574 | -4.3 | 0.048 | 0.18 | -2.44 |
| 993.592 | -6.9 | 0.017 | 0.17 | -2.59 |
| 994.598 | -6.4 | 0.021 | 0.18 | -2.50 |
| 1013.31 | -13.3 | 0.012 | 0.02 | -5.57 |
| 1164.36 | -11.7 | 0.012 | 0.04 | -4.79 |
| 1165.36 | -8.9 | 0.013 | 0.05 | -4.42 |
| 1168.66 | -4.7 | 0.037 | 0.33 | -1.61 |
| 1170.69 | -5.6 | 0.027 | 0.06 | -3.95 |
| 1182.62 | -5.5 | 0.027 | 0.09 | -3.48 |
| 1208.35 | -10.0 | 0.012 | 0.02 | -5.52 |
| 1209.36 | -11.2 | 0.012 | 0.03 | -5.17 |
| 1214.63 | -5.1 | 0.032 | 0.09 | -3.43 |

*A fold change (FC) > 1 indicate that the relative intensity of this metabolite is higher in B1-anemones, whereas a FC < 1 indicates that it is higher in $C1^{acro}$-anemones. For instance, a FC value of 0.1 means that the relative intensity of this metabolite is 10 times higher in $C1^{acro}$-anemones than B1-anemones.

**Table S6. Saturation state of the betaine lipids**

| m/z | Annotation | Saturation | Possible structure |
|---|---|---|---|
| 472.363 | DGTS | Polyunsaturated | 15:2 |
| 490.374 | MGCC | Saturated | 16:0 |
| 512.359 | MGCC | Polyunsaturated | 18:3 |
| 562.374 | MGCC | Highly polyunsaturated | 22:6 |
| 750.589 | DGCC | Saturated | 32:0 |
| 763.512 | DGCC | Highly polyunsaturated | 18:4/16:4 |
| 772.572 | DGCC | Polyunsaturated | 36:6 |
| 774.589 | DGCC | Polyunsaturated | 36:5 |
| 798.587 | DGCC | Monounsaturated/polyunsaturated/ highly polyunsaturated | 38:7/36:4 |
| 800.604 | DGCC | Polyunsaturated/ highly polyunsaturated | 38:6 |
| 814.619 | DGCC | Monounsaturated/ Polyunsaturated | 38:1/39:6 |
| 822.586 | DGCC | Polyunsaturated/ highly polyunsaturated | 40:9/38:6 |
| 823.588 | DGCC | Polyunsaturated/ highly polyunsaturated | 40:9/38:6 |

**Table S7. Details of Symbiodiniaceae culture used to inoculate the anemones *E. diaphana*.**

| Species | ITS2 | Details | Culture ID | °C | Original host | Host origin |
|---|---|---|---|---|---|---|
| *Breviolum minutum* | B1 | Homologous, wild-type | SCF 127-01 | 27 | *E. diaphana* | Central GBR, Australia |
| *Cladocopium* C1$^{acro}$ | C1 | Heterologous, wild-type | SCF 055-01.10 | 27 | *Acropora tenuis* | Magnetic Island, Australia |
